# GET: a foundation model of transcription across human cell types

**DOI:** 10.1101/2023.09.24.559168

**Authors:** Xi Fu, Shentong Mo, Alejandro Buendia, Anouchka Laurent, Anqi Shao, Maria del Mar Alvarez-Torres, Tianji Yu, Jimin Tan, Jiayu Su, Romella Sagatelian, Adolfo A. Ferrando, Alberto Ciccia, Yanyan Lan, David M. Owens, Teresa Palomero, Eric P. Xing, Raul Rabadan

## Abstract

Transcriptional regulation, involving the complex interplay between regulatory sequences and proteins, directs all biological processes. Computational models of transcription lack generalizability to accurately extrapolate in unseen cell types and conditions. Here, we introduce GET, an interpretable foundation model designed to uncover regulatory grammars across 213 human fetal and adult cell types. Relying exclusively on chromatin accessibility data and sequence information, GET achieves experimental-level accuracy in predicting gene expression even in previously unseen cell types. GET showcases remarkable adaptability across new sequencing platforms and assays, enabling regulatory inference across a broad range of cell types and conditions, and uncovering universal and cell type specific transcription factor interaction networks. We evaluated its performance on prediction of regulatory activity, inference of regulatory elements and regulators, and identification of physical interactions between transcription factors. Specifically, we show GET outperforms current models in predicting lentivirus-based massive parallel reporter assay readout with reduced input data. In fetal erythroblasts, we identify distal (>1Mbp) regulatory regions that were missed by previous models. In B cells, we identified a lymphocyte-specific transcription factor-transcription factor interaction that explains the functional significance of a leukemia-risk predisposing germline mutation. In sum, we provide a generalizable and accurate model for transcription together with catalogs of gene regulation and transcription factor interactions, all with cell type specificity.

## Main

Transcriptional regulation constitutes a critical yet largely unresolved domain, underpinning diverse biological processes, including those associated with human genetic diseases and cancers^1^. A conserved regulatory machinery orchestrates transcriptional changes, including transcription factors that bind to regulatory sequences; coactivators, mediators, and core transcriptional factors; and RNA Polymerase II^2–4^. While different cell types may possess different subsets of regulatory regions, the biochemistry of protein-protein interaction and protein-DNA interaction remains largely the same across cell types when epigenetic conditions are fixed. Clustering of known transcription factor binding site motifs^5^ demonstrates great functional redundancy in transcription factor DNA binding domains, further reducing the combinatorial variability of regulatory interactions. However, our understanding of transcription regulation is often limited to specific cell types, and it is not clear how the combinatorial interaction of different transcription factors determines the diversity of expression profiles observed across cell types.

Advances in sequencing technology and the adoption of sophisticated machine learning architectures have enabled the exploration of expression and associated noncoding regulatory features across a broad spectrum of cell types. Traditional methods such as Expecto^6^ and Basenji2^7^ utilized convolutional neural networks for shorter input sequences, while state-of-the-art approaches like Enformer^8^ extended capabilities with the transformer architecture. Nonetheless, existing models present challenges. A key limitation is that they can only make predictions on the training cell types, hindering the generalizability and utility of the model.

In the landscape of machine learning and computational biology, foundation models like GPT4^9^ and ESM2^10^ are emerging as a transformative approach. These models serve as a foundation, upon which specialized adaptations can be built to address specific tasks or challenges. By utilizing extensive pretraining on broad and diverse datasets, foundation models provide a generalized understanding of underlying patterns and relationships. In the field of transcriptional regulation, a foundation model has the potential to synthesize the vast complexities of regulatory mechanisms across various cell types, offering a versatile framework that can be finetuned to target specific applications, cell types, or conditions. For example, recent developments in single cell foundation models like Geneformer^11^, scGPT^12^, and scFoundation^13^ have demonstrated how diverse transcriptome profiles can be encoded in one model, enabling various contextualized downstream tasks including cell type annotation and perturbation prediction. However, a foundation model of how transcription emerges from the chromatin landscape has not yet been explored.

Here we introduce the general expression transformer (GET, **Supplementary Figure 1**), an interpretable foundation model for transcriptional regulation across 213 human fetal and adult cell types that exhibits universal applicability and exceptional accuracy. GET learns transcriptional regulatory syntax from chromatin accessibility data across hundreds of diverse cell types and successfully predicts gene expression in both seen and unseen cell types, approaching experimental accuracy (**Figure 1a, Supplementary Figure 2a**). The versatile nature of GET allows it to be transferred to different sequencing platforms and measurement techniques. Additionally, it offers zero-shot prediction of reporter assay readout in new cell types, potentiating itself as a prescreening tool for cell type specific regulatory elements. GET outperforms previous state-of-the-art models in identifying cis-regulatory elements, and identifies novel and known upstream regulators of fetal hemoglobin. Through interpreting GET, we provide rich regulatory insight for almost every gene in 213 cell types. Using coregulation information predicted by GET, we performed causal discovery to pinpoint potential motif-motif interactions and constructed a structural interaction catalog of human transcription factors and coactivators. Using information provided by GET, we successfully identified a lymphocyte-specific transcription factor-transcription factor interaction involving PAX5 and nuclear receptor family transcription factors, and highlighted a possible disease driving mechanism of a leukemia-associated germline variant which affects the binding of the PAX5 disordered region to the nuclear receptor domain of retinoic acid receptors. Overall, GET’s broad applicability and profound understanding of transcriptional regulation will advance understanding of noncoding genetic variants and guide de novo design of cell-type specific transcriptional regulatory circuits and transcription factors for synthetic biology applications.

**Figure 1.**
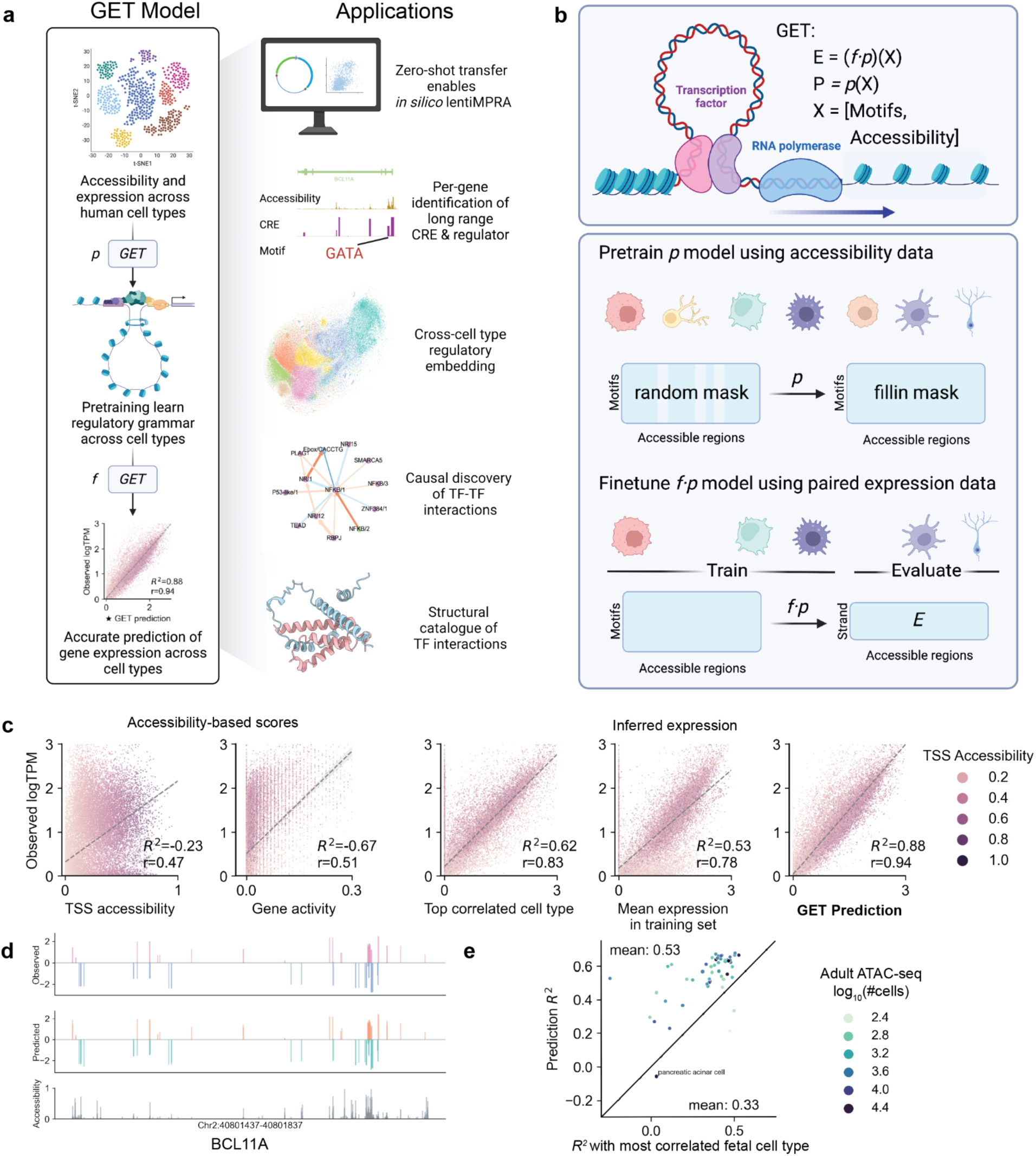
The GET model’s universal applicability and exceptional accuracy as a transcription foundation model. **a**. GET derives transcriptional regulatory syntax (<u>p</u>retrain) from chromatin accessibility data across hundreds of cell types, providing reliable predictions (finetune) of gene expression in both seen and unseen cell types. The model’s broad applicability and comprehensibility allow for zero-shot prediction of lentiMPRA measurements, extensive identification of cell-type-specific regulatory elements and upstream transcription factors, universal embeddings of regulatory information, and causal understanding of transcription factor-transcription factor interactions. **b**. Schematic illustration of the training scheme of GET. The input of GET is a peak (accessible region) by transcription factor (motif) matrix derived from a human single cell (sc) ATAC-seq atlas, summarizing regulatory sequence information across a genomic locus of more than 2 Mbps. Through self-supervised random masked pretraining of the input data across more than 200 cell types, GET learns transcriptional regulatory syntax (***p***). Finetuned on paired single cell ATAC-seq and RNA-seq data, GET learns to transform the regulatory syntax to gene expression even in leave-out cell types (**f⋅p**). **c**. Benchmark of GET prediction performance on unseen cell types (Fetal astrocyte). Each point is a gene. Color represents normalized chromatin accessibility in TSS. Gene activity is a score widely used in modern scATAC-seq analysis pipelines^21^. Top correlated cell type is the training cell type whose observed gene expression has the largest correlation with Fetal astrocyte, in this case Fetal inhibitory neuron. Mean cell type is the mean observed gene expression across training cell types. Dashed line represents linear fits. **d.** Example visualization of observed expression (top, log_10_TPM), GET prediction (mid, log_10_TPM) and chromatin accessibility (bottom, log_10_CPM) of the BCL11A locus in Fetal erythroblast. Positive (negative) values represent expression on positive (negative) strand on hg38. **e**. GET trained on fetal cell types generalizes to adult cell types without retraining, outperforming the most correlated cell type baseline. X axis showing R^2^ score between GET prediction in adult cell types and observed expression in most similar fetal cell types. Y axis showing R^2^ score between GET prediction and observed expression in the adult cell type.

### GET, a foundation model for transcription regulation across 213 human cell types

We embarked on developing GET, a novel foundational model to comprehend transcriptional regulation across a diverse range of cell types. Unlike previous models such as Enformer^8^, GET employs an extensive effective sequence length exceeding 2 Mbps (**Method: Computational cost comparison**) and is not constrained to making predictions in only training cell types.

The design philosophy of GET is rooted in the conceptual understanding of transcription regulation (**Figure 1b**). At the forefront, Promoter and related contextual regulatory elements can be characterized by how well they bind different transcription factors (motif binding score, **Methods-Motif features**) and how accessible they are in specific cell types. These features shape a chromatin environment (*p(X)*) that governs how RNA polymerase II (PolII) functions. Using an embedding and attention architecture^14^ specifically designed for the regulatory elements (**Methods-Model architecture**), we performed self-supervised pretraining to allow GET to learn how the regions and features interact with each other across diverse cell types. Specifically, by randomly masking out regulatory elements, the model is trained to predict the motif binding scores and optionally accessibility score in the masked region. Subsequently, PolII will readout the chromatin environment *p(X)* into an expression *E* (**Figure 1b**). A finetuning stage with the same architecture but a different output head will simulate this process. This two-stage design makes it possible to use chromatin accessibility data with no paired expression measurement, greatly improving the diversity of regulation information in the training data.

The pretraining of GET uses pseudobulk chromatin accessibility gathered from single cell assay for transposase-accessible chromatin with sequencing (scATAC-seq) data across 213 human fetal and adult cell types^15–17^. Out of these, 153 were coupled with expression data, acquired either through a multiome protocol or separate single cell RNA sequencing (scRNA-seq) experiments^18,19^ (**Methods: ATAC-seq data processing** and **RNA-seq data processing**). We calculated the motif binding score using known position weight matrices and summarized them according to sequence similarity to reduce feature redundancy^5^. Assuming additivity in motif binding score, every sample is a region-by-feature matrix derived from a continuous range on the accessible genome across different cell types. This design of model input ensures both cell type specificity and generalizability while enabling efficient computational modeling. Strand-specific expression values are assigned to each region based on their overlap with an expressed gene’s promoters.

### GET accurately predicts gene expression in unseen cell types at experimental accuracy

We first assessed GET’s ability to accurately predict gene expression in unseen cell types in a setting where one cell type is left out during the expression finetuning process. Remarkably, GET demonstrated the capacity to consistently predict the expression of the left-out cell types at a level of accuracy comparable to experimental standards, even when trained without quantitative accessibility signals. An example can be taken from left-out astrocytes, where the Pearson correlation between GET’s predicted expression values and the observed expression reached 0.94 (R² = 0.88) (**Figure 1c,d**), a result that is in line with experimental accuracy across different culture systems and biological replicates of human astrocytes^20^ (Pearson r = 0.92-0.99, **Supplementary Figure 2a**). GET’s performance surpasses both transcription start site (TSS) accessibility (r = 0.47, R² = -0.23) and gene activity score^21^ (r = 0.51, R² = -0.67), emphasizing the significance of DNA sequence specificity and distal context information in transcription regulation. Furthermore, GET managed to outperform two robust benchmarks, including top correlated cell type expression (r = 0.83, R² = 0.62) and mean expression across training cell types (r = 0.78, R² = 0.53; as illustrated in **Figure 1c**). Additional validation was carried out which confirmed GET’s capability to make cell-type-specific predictions, as evidenced by a Pearson correlation of 0.91 (R² = 0.82) between predicted and observed log fold change for Fetal astrocyte and Fetal erythroblast expression (see **Supplementary Figure 2b**). GET surpasses the mean expression baseline across all evaluated fetal cell types as well as Enformer’s performance via linear probing from its CAGE output tracks on the same set of cell types (**Supplementary Figure 2c**).

We proceeded to investigate GET’s generalizability to adult cell types when trained solely on fetal data. Our findings showed an average R² of 0.53 across diverse adult cell types, once again surpassing the baseline (R² = 0.33) obtained using corresponding fetal cell types for prediction (e.g., utilizing Fetal astrocyte to predict Adult astrocyte) (**Figure 1e**). The only 3 cell types where we cannot beat the baseline are cell types with low cell counts in either ATAC-seq (Blood Brain Barrier Endothelial Cell and Tuft Cell, n<600) or RNA-seq label (pancreatic acinar cell). This result reinforces the proposition that GET can extract common regulatory mechanisms that span various cell types and stages of life.

To ascertain the impact of cross-cell-type pretraining on prediction performance, we finetuned a GET model from random initialization, which exhibited a substantial drop in performance compared to the pretrained version with the same number of training epochs (Pearson r: 0.596; Spearman rho: 0.642, **Supplementary Figure 2d**). Extending the training period for this baseline to 800 epochs failed to enhance its performance (Pearson r: 0.607; Spearman rho: 0.658, **Supplementary Figure 2d**), highlighting the essential role of pretraining in facilitating model generalization. GET also shows superior performance when compared against common supervised machine learning approaches on expression prediction (**Supplementary Figure 2e**). When finetune GET just one sample using a leave-one-chromosome-out setting, we get consistently performance across chromosomes (Mean Pearson r: Fetal Astrocyte: 0.773; **Supplementary Figure 2f**)

In summation, our study demonstrates that by leveraging widely accessible ATAC-seq data and established transcription factor binding motifs, GET acquires a broad understanding of the regulatory code, empowering it to predict unseen cell type expression with experimental precision.

### GET can be transferred to different sequencing platforms and non-physiological cell types

Diverse data generation platforms and processing methods often present a significant challenge for the universal generalizability of pretrained models. To assess whether GET can be transferred to new sequencing platforms in such scenarios, we benchmarked its performance on 10x multiome sequencing of the lymph node^22^ (**Figure 2a, Methods: ATAC-seq data processing** and **RNA-seq data processing**) using the leave-cell-type-out evaluation approach. Notably, GET maintained consistent prediction outcomes for both finetuned or left-out cell types (**Figure 2b**).

**Figure 2.**
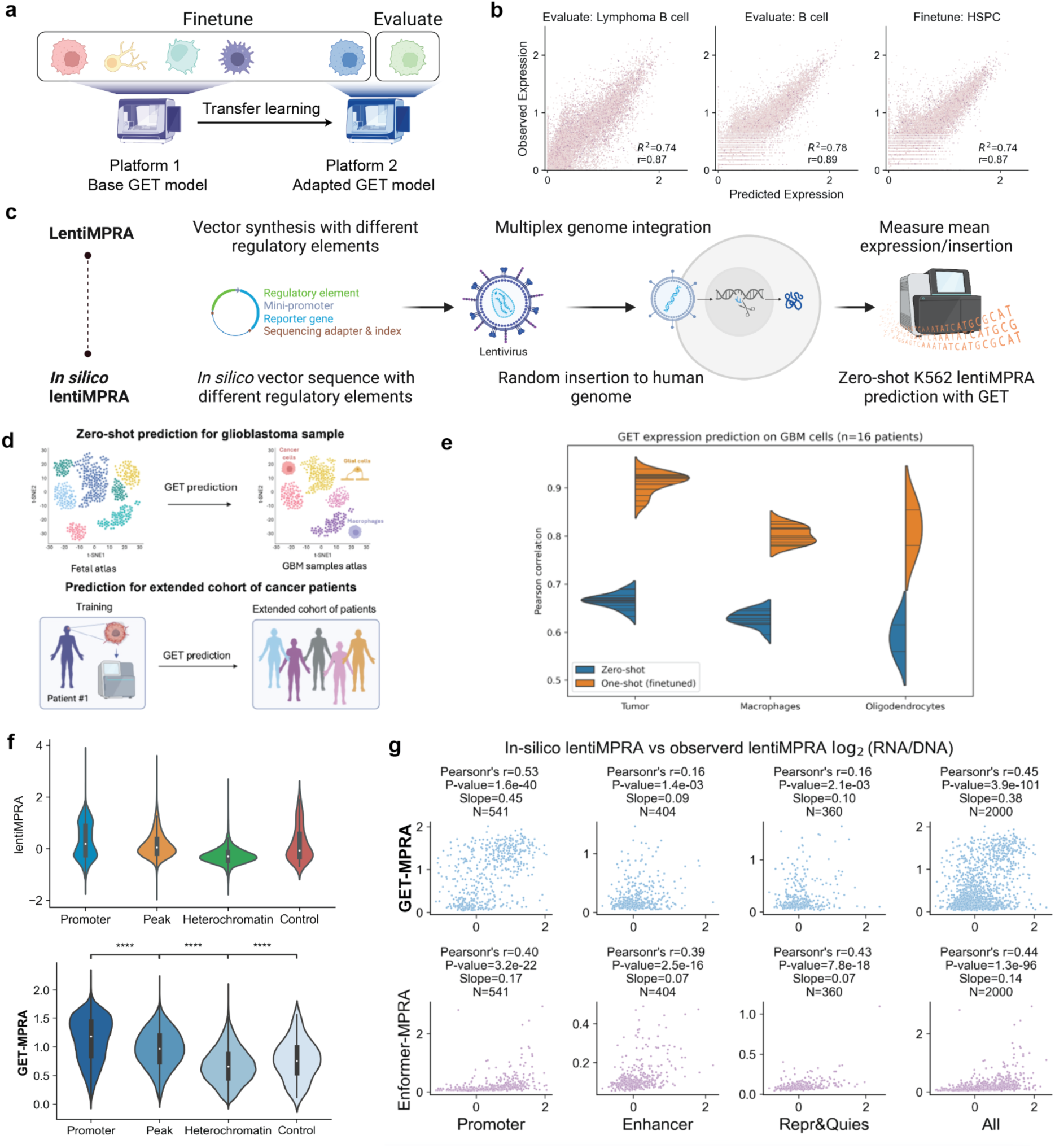
Transfer learning adapts GET to new platforms and measurements. **a**. Schematic illustration of transferring GET to a lymph node 10x multiome dataset. **b**. Finetuned GET accurately predicts expression in training and leave-out evaluation cell types. **c**. Schematic workflow of lentiMPRA experiments and *in silico* lentiMPRA using GET model finetuned on K562 multiome data. **d.** Schematic showing the application of GET in the zero-shot setting to predict gene expression from GBM patient samples (top) and finetuned on a single GBM patient sample used to predict gene expression for an extended cohort of GBM patients. **e.** Pearson correlation scores for GET expression prediction on GBM cells (n = 16 samples) comparing tumor, macrophages, and oligodendrocytes for zero-shot and one-shot (finetuned). **f.** Readout distribution of lentiMPRA (log_2_RNA/DNA, top) and GET prediction (mean expression across genomic insertions, bottom) for different types of elements. Two sided Mann-Whitney U-test: Promoter vs. Peak: p < 1e-237; Peak vs. Heterochromatin: p < 1e-237; Heterochromatin vs. Control: p = 4.049e-04. ***: 1.00e-04 < p <= 1.00e-03; ****: p <= 1.00e-04. **g.** Benchmark of GET lentiMPRA prediction against Enformer on random subset of elements. X axes show observed lentiMPRA readout (log_2_RNA/DNA). Y axes show predicted expression in log_10_ TPM.

We also analyzed the potential of the GET model to investigate non-physiological cell types, including patient tumor cells. We applied GET to 10x multiome data from 18 glioblastoma (GBM) IDH1 wild-type patients (**Figure 2c**)^23^. The zero-shot performance on GBM tumor cells across held-out patients achieved a mean Pearson correlation of 0.68. When finetuning GET on a single patient sample, the Pearson correlation improved to above 0.90 (**Figure 2d**). Leave-one-chromosome-out finetuning on a single patient showed a mean Pearson of 0.73 across chromosomes (**Supplementary Figure 2f**).

Given GET’s demonstrated adaptability, we explored its applicability to other experimental assays. We applied GET to predict cap analysis of gene expression (CAGE) from the K562 cell line as a test of sequencing platform shift. When comparing finetuned GET to Enformer’s predictions for K562 CAGE, GET achieved a Pearson correlation of 0.90 compared to Enformer’s performance of 0.74 (**Supplementary Figure 2g**).

We finetuned GET on scATAC-seq from fetal astrocytes, finding a Pearson correlation of 0.71 comparable to Enformer’s prediction of DNase for the same cell type (**Supplementary Figure 2h**). Using only motif inputs, we finetuned GET to predict quantitative K562 peak-level chromatin accessibility from bulk ENCODE OmniATAC^24^ data, achieving a 0.81 Pearson correlation across different leave-one-chromosome-out experiments (**Supplementary Figure 2h**, middle, N=22). This is not directly comparable to sequence-based accessibility models like ChromBPNet^25,26^ since the variable peak sequence length will be partially encoded in the motif binding score. As a comparison, GET finetuned on fixed-length fetal astrocyte scATAC data resulted in a 0.72 Pearson correlation. We also found these finetuned ATAC-prediction models are relatively robust to leave-out-motif training when the number of omitted motifs is less than 10, indicating that the model benefits from the masked prediction objective and shared sequence information between motifs (**Supplementary Figure 2h**, right, **Method: Leave-out-motif evaluation**).

### Zero-shot GET prediction of expression-driving regulatory elements in new cell types

Building on the versatility of GET across diverse platforms and measurements, we ventured into examining its capacity for zero-shot prediction of expression-driving regulatory elements in unseen cell types. Lentivirus-based massively parallel reporter assay (lentiMPRA) provides a robust mechanism to test the regulatory activity of numerous genetic sequences by integrating them into the genome, thereby circumventing the limitations inherent in episomal MPRAs and ensuring relevant biological readouts in hard-to-transfect cell lines^27^. Recently utilized to assess over 200,000 sequences in the K562 cell line, this experimental assay has created a comprehensive benchmark dataset for evaluating whether the GET model can identify regulatory elements in a cell-type specific context^28^ (**Figure 2e**).

In an *in silico* procedure akin to the lentiMPRA experiment, we employed the GET model finetuned on bulk ENCODE K562 OmniATAC chromatin accessibility and expression data from NEAT-seq. We then constructed the sequences for insertion, including both the regulatory sequence and minipromoters, randomly inserting these sequences across the genome. Utilizing the GET model, which was not trained on lentiMPRA data, we inferred the activity of the mini promoter within the corresponding chromatin context and averaged over all insertions to obtain a mean readout indicative of the regulatory activity (**Figure 2e, Methods:LentiMPRA zero-shot prediction**). We found that optimal performance is achieved when we combine the mean predicted expression readout with mean predicted accessibility of the inserted element (**Supplementary Figure 3a-b**).

Upon examination of the readout distribution for different types of elements (**Figure 2f**), we found our predictions to be consistent with experimental data. Promoter sequences exhibited the highest GET-MPRA readout, followed by chromatin accessibility peak sequences. Heterochromatin sequences registered the lowest readouts, and control sequences spanned a wide range of readout values.

When benchmarking our model against Enformer, which was trained on 486 tracks of functional K562 genomics data, including transcription factor and histone modifications chromatin co-immunoprecipitation sequencing (ChIP-seq), cap analysis of gene expression (CAGE), and chromatin accessibility measurements, we discovered that our model made more accurate predictions overall (Pearson’s r = 0.56 versus 0.44, see **Figure 2g**). Enformer outperformed in peak regulatory activity predictions, which can be attributed to its nucleotide-level modeling architecture and extensive training data specifically targeting K562. Overall, Enformer’s predictions tend to have larger across-genome variance (**Supplementary Figure 3c**). GET also presented significant advantages in computational cost. In fact, for this comparison, we had to subsample to 2,000 elements to complete the calculation with Enformer in 3 days. While using the same amount of computing time GET can screen all 200,000 elements (see **Methods:Computational cost comparison**).

### GET accurately identifies cis-regulatory elements and upstream regulators

Single-cell multiome studies enable the identification of cis-regulatory elements (CRE) in specific cell types, offering potential phenotype intervention targets. Traditional peak-to-gene workflows largely depend on correlating multiome ATAC-seq and RNA-seq counts, with regulator identification necessitating additional filtering by transcription factor (TF) expression^29–31^. Such methods are limited in comprehensiveness due to the nonlinear relationship between accessibility and transcription level and sparsity of single cell data and can usually can only produce results for thousands of genes. Through model interpretation techniques (**Methods:Model interpretation**), we can efficiently derive region or motif contribution scores for expressed genes across cell types, producing results for virtually all genes in even less abundant cell types (approximately 1,000 cells). Focusing on fetal erythroblasts, we leveraged published genome base-editing data to investigate four known fetal hemoglobin regulating loci^32^ (*BCL11A, NFIX, KLF1, HBG2*, where the first three are known to regulate fetal hemoglobin and *HBG2* encodes a fetal hemoglobin subunit, **Figure 3a**).

**Figure 3.**
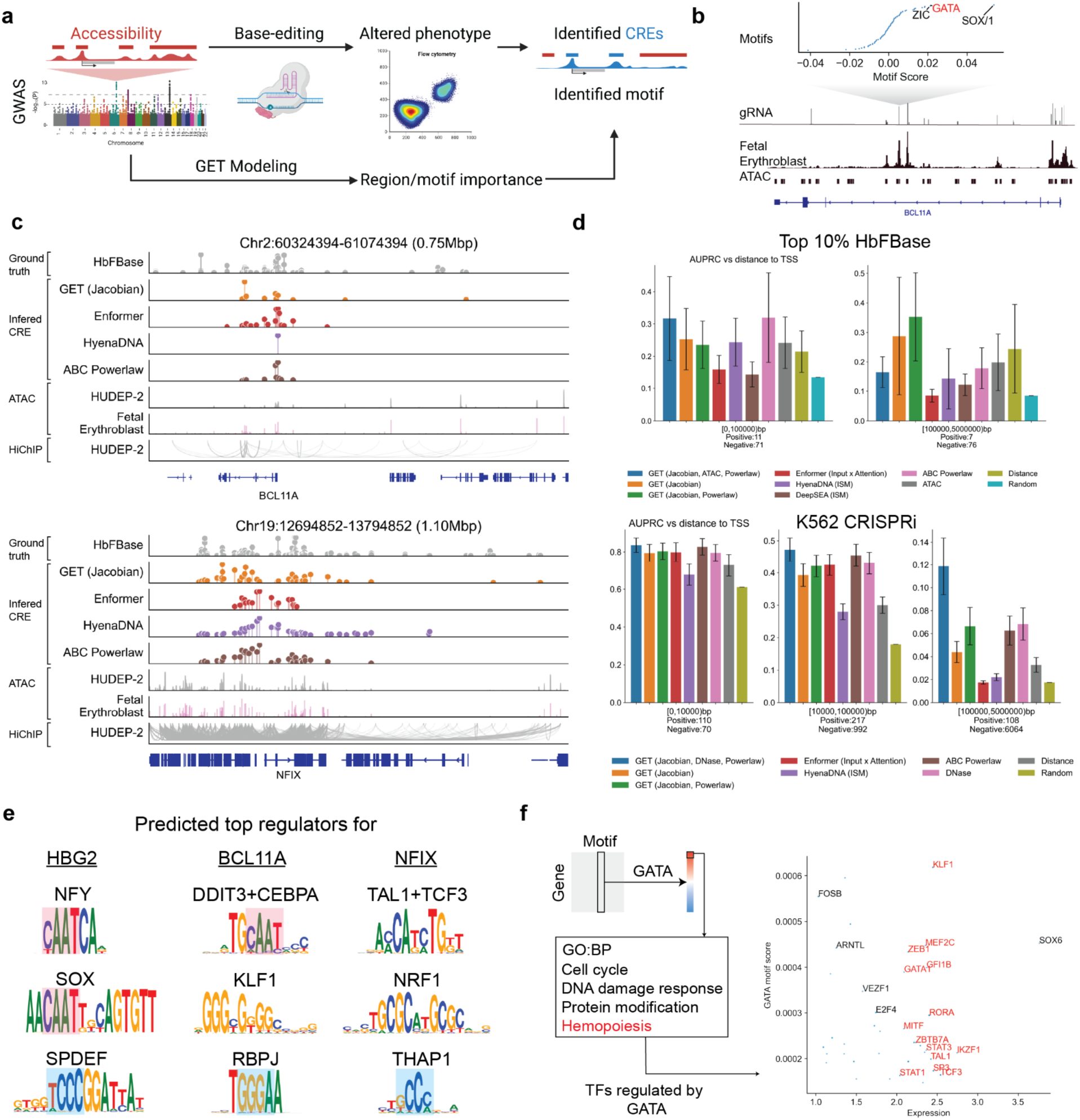
The GET model identifies cell-type-specific regulator and cis-regulatory elements. **a.** Case study of identifying cis-regulatory elements (CRE) and regulators controlling a phenotype, fetal hemoglobin (HbF) level. Four genome-wide association loci (BCL11A, MYB, NFIX, and HBG2) have been subjected to genome editing in a previous study, providing the labels for GET benchmarking. Region/motif contribution score for each gene can be computed using the GET model. **b.** GET identifies the GATA motif in erythroid-specific enhancer that upregulates BCL11A, an HbF repressor. (Top) motif contribution score for BCL11A expression in the erythroid-specific enhancer. (Mid) gRNA enrichment score (HbFBase). Higher score means enrichment in high HbF cells, which implies these edits disturb a cis-regulatory element or regulator binding site that can upregulate BCL11A. (Bottom) single cell ATAC-seq signal and peak from Fetal erythroblast. **c**. Genome tracks displaying inferred cis-regulatory elements (CREs) for BCL11A and NFIX loci. Plots for HBG2 and MYB loci can be found in **Supplementary Figure 4c**. The loci shown are Chr2:60324394-61074394 (0.75 Mbp) and Chr19:12694852-13794852 (1.10 Mbp). From top to bottom, the tracks represent: HbFBase, showing the gRNA enrichment score from base-editing experiments; GET, showing the inferred region importance score; Enformer, showing the inferred region importance score; HyenaDNA, showing the *in silico* mutagenesis (ISM) result using the pretrained HyenaDNA language model; ABC Powerlaw, showing the Activity-by-Contact prediction using fetal erythroblast ATAC and K562 Hi-C power law; ATAC-seq data from HUDEP-2, an erythroblast cell line; ATAC-seq data from fetal erythroblast cells, used in the training of GET; and HiChIP-seq data from HUDEP-2, demonstrating chromatin interactions. **d**. Benchmark results comparing GET to other methods for predicting enhancer-promoter pairs, including analysis of distal (>100kb) interactions. (Top) Erythroblast fetal hemoglobin regulating enhancer prediction. (Bottom) K562 CRISPRi enhancer target prediction. Area under precision-recall curve (AUPRC) is shown. Ablation of different GET prediction components (Jacobian, DNase, Powerlaw; see **Method: Predict enhancer targets**) is also shown in the plot. **e**. Predicted top three regulators (motifs) for BCL11A, NFIX, and HBG2. Similar sequence patterns are highlighted with color shades. **f**. GATA downstream targets inferred by GET (top 10% motif score) show functional enrichment in hemopoiesis. Scatterplot shows predicted gene expression (X-axis) and GATA-motif score (Y-axis) for GATA-targeted genes with predicted expression larger than 1. All transcription factors among these genes are labeled in the plot, where those involved in Hemopoiesis are highlighted in red.

Applying GET to fetal erythroblasts yielded interesting insights into the regulation of fetal hemoglobin. We rediscovered the central role of the GATA transcription factor, which, via its binding to an erythroid-specific enhancer, orchestrates the expression of BCL11A, a known modulator of hemoglobin regulation^33,34^. Interestingly, GET also highlighted the role of the SOX family of transcription factors in this enhancer, which were previously linked to fetal hemoglobin^35^ but not known to function through this specific enhancer (**Figure 3b**).

Examining all four loci—BCL11A, NFIX, KLF1, and HBG2—we benchmarked GET against established models like Enformer^8^, HyenaDNA^36^, DeepSEA^37^, and Activity-by-Contact (ABC)^38,39^ . Distinctly, GET outperformed these counterparts, especially in detecting long-range enhancer-promoter interactions (**Figure 3c-d, Supplementary Figure 4a-c**). We also show that while enhancer chromatin accessibility is predictive of regulatory activity for proximal enhancer-promoter relationships, its precision diminishes for long-range interactions. Alternative evaluations using different functional enhancer thresholds (top 10% or 25% of the experimental readout, HbFBase) reaffirm GET’s precision in this scenario (**Figure 3d top**). We also benchmarked the same task for the K562 cell line using a CRISPRi based benchmark dataset from ENCODE^40^, finding a similar conclusion (**Figure 3d bottom, Supplementary Figure 4d**).

GET is able to extract overall motif importance across cis-regulatory elements (CREs) for specific genes. For HBG2, BCL11A, and NFIX, the top motifs identified were consistent with their known transcriptional regulators or hematopoietic transcription factors (**Figure 3e**). For instance, we found significance of NFY and SOX motifs for HBG2 and the reaffirmation of KLF1’s influence on BCL11A^32^. Additionally, for NFIX, GET adeptly pinpointed the involvement of TAL1, a known GATA1 binding partner and hematopoietic factor^41^.

To determine downstream targets for specific regulators, we developed an *in silico* analysis, taking the GATA motif as a case study. Using the GET motif contribution matrix, we spotlighted the top 10% of genes influenced by the GATA motif. Notably, aligning with GATA1’s status as a master regulator of erythroid development, the hematopoiesis biological process was enriched^42^ (Fisher exact test, multiple hypothesis adjusted P-value=7.6×10^−4^, **Figure 3f** and **Methods:Gene ontology enrichment of top target genes of a regulator**) within this gene set. Known erythroid lineage transcription factors like KLF1, GATA1, TAL1, and IKZF1 were predicted to be regulated by the GATA motif, underscoring GATA’s pivotal role in a multifaceted regulatory network, in line with existing literature^43^.

To assess GET’s capability to detect significant regulatory alterations across different cell states, we focused on the differential expression between fetal erythroblast and fetal hematopoietic stem cells (HSC). Our expectation is that the genes that mark the lineage differentiation should show larger gradients from lineage specific transcription factors than those that are indifferent across lineages. Our findings confirmed this, as we noted substantial Spearman correlation between the motif contributions in erythroblasts and the differential expression log fold changes for several erythroblast-related transcription factors, including GATA, ZBTB7A, and MZF1 (**Supplementary Figure 4e**).

Extending our analysis to encompass all fetal and adult cell types, we found that for certain well-known regulators, such as CTCF, MBD2, NFKB, and NFI, there exists a significant correlation between the mean expression of inferred target genes and the mean expression of all regulators within the corresponding family (e.g., NFIA, NFIB, NFIC, NFIX for the NFI motif, **Supplementary Figure 5a**). Overall, we conclude that GET possesses the ability to learn meaningful regulatory information that is naturally transferable between cell states.

### Cell-type specific regulatory insights through cross-cell-type embedding with GET

Utilizing a cross-cell-type architecture, GET is configured to extract the regulatory context for genes spanning various cell types, embedding them within a shared high-dimensional space (**Methods:Regulatory embedding**). In order to further visualize what GET learns from different cell types, we explored different embedding layers of GET. We found that the embedding from final layers correlates well with expression levels, while earlier layers are more indicative of differences in regulatory grammar (**Supplementary Figure 5b**).

To investigate whether the embedding tied to regulatory grammar retains cell type-specific information, we gathered the first transformer layer’s embedding for all promoters across cell types. This allows us to capture not only the motif information of the promoters but also that of the CREs owing to the attention mechanism. Intriguingly, a t-distributed stochastic neighbor embedding (t-SNE)^44^ visualization of randomly sampled embeddings showed motif separation but no cell type differentiation, suggesting that with a sufficient number of cell types, most regulatory grammar is shared across cell types, although they may be instantiated on different genes (**Figure 4a**).

**Figure 4.**
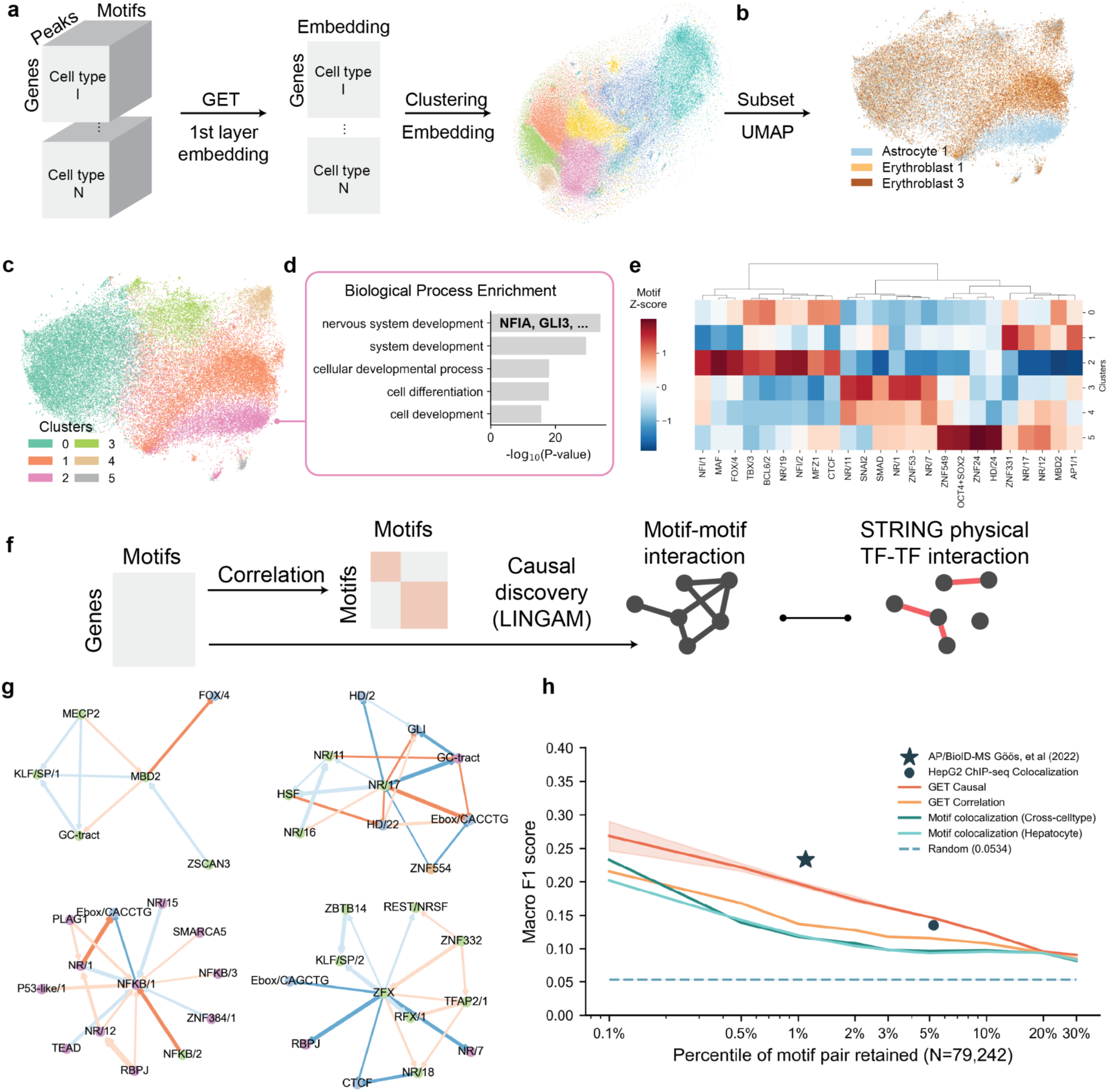
GET captures regulatory information across cell types and informs casual transcription factor-transcription factor interaction. **a**. Workflow to collect and visualize cross-cell-type regulatory embedding, showing a tSNE visualization of the resulting embedding space colored by Louvain clustering. **b**. The cross-cell-type regulatory embedding reveals cell-type specificity in transcriptional regulation. Subsampled embedding from Fetal astrocyte (blue) and two Fetal erythroblast (yellow and brown) cell types are visualized with UMAP. **c**. Louvain clustering of subsampled embedding in panel b. Note that cluster 2 is an astrocyte specific cluster. **d**. Gene ontology enrichment of genes in cluster 2, showing astrocyte-relevant terms and astrocyte marker genes e.g. NFIA, GLI3. **e**. GET motif contribution Z-score (red means higher score compared to other clusters) for each cluster. Note that cluster 2 has elevated NFI/1 and NFI/2 motifs, which correspond to the NFI family transcription factors. **f**. Causal discovery using the GET motif contribution matrix identifies transcription factor-transcription factor interactions. Physical interactions from STRING databases are used as a benchmark to calculate the concordance. **g**. Example causal neighbor graph showing interactions (edges) between motifs (nodes). Edge weights represent interaction effect size. Edge directions mark casual directions. Blue and red edge color marks negative or positive estimated causal effect size by LiNGAM, respectively. Node color marks community detected on the full causal graph. In-community edges are marked by reduced saturation. **h**. Benchmark of concordance of inferred transcription factor-transcription factor interactions using different methods with physical interactions from the STRING database. X-axis marks different cutoffs of retained interaction in percentile of 79,242 total possible interactions. Y-axis marks the ratio of selected interactions that is also marked as interacted in STRING. Green line marks the random selection background. Orange line marks the result of selection using motif-motif contribution score correlation. Red line marks the causal discovery result. Shaded area marks standard error across 5 bootstraps. Green and aqua lines show results from motif colocalization, computed as correlation between motif binding vectors in accessible regions across all cell types (green) or in hepatocytes (aqua). The star marks the result from a recent mass-spectrometry-based transcription factor-transcription factor interaction atlas^57^ (0.23 Macro F1 at 1.09% recall). The round dot marks the performance of a colocalization score computed from 677 HepG2 TF ChIP-seq (0.13 Macro F1 at 5.24% recall, **Method:Causal discovery of regulator interaction**).

Nonetheless, when we subset the embeddings to only three specific cell types (fetal astrocyte and two fetal erythroblast subclusters), the UMAP^45^ exhibited distinct clusters for astrocytes and erythroblast genes (**Figure 4b**). This result further corroborates that GET is proficient at discerning cell-type-specific regulatory information.

Looking further into the astrocyte-specific gene cluster (cluster #2 in **Figure 4c**), we discovered that this gene set is particularly enriched in the development of the nervous system and includes astrocyte transcription factors such as NFIA^46,46,47^ and GLI3^48^ (**Figure 4d**). Moreover, a comparison of motif contribution across clusters revealed a higher presence of NFI motifs in the astrocyte-specific cluster (**Figure 4e**), shedding light on the unique regulatory program within astrocytes.

### GET-based causal discovery identifies potential transcription factor-transcription factor interactions

Given GET’s proficiency in elucidating regulatory mechanisms across diverse cellular contexts, we next investigate whether it learns transcription factor-transcription factor functional interactions implicitly (**Figure 4f**). Using a cell-type agnostic gene-by-motif matrix (**Methods:Causal discovery of regulator interaction**), we evaluated the correlation between different motif vectors (**Supplementary Figure 6a**). High correlation may represent common genomic targets between different transcription factors. Transcription factor pairs with correlation values in the top decile are more likely to participate in the same biological functions compared to those in the bottom decile (**Supplementary Figure 6b**, Kolmogorov–Smirnov test, P-value=6.78×10^−82^). For example, MBD2 and MECP2, a high correlation transcription factor pair, both act as readers of DNA methylation^49,50^.

We further extended our investigation of motif-motif interactions by utilizing a causal discovery algorithm, Linear Non-Gaussian Acyclic Model (LiNGAM)^51^, to derive a directed acyclic graph from the cell-type agnostic gene-by-motif matrix (**Methods:Causal discovery of regulator interaction**). The consequent network, displaying interactions with an absolute value greater than 0.1 for clarity in visualization, are shown in **Supplementary Figure 6c**. We identified factors such as CTCF, KLF/SP/2 (GC rich motif), TFAP2/1, ZFX, RBPJ, Accessibility, and methylation-associated E2F as having the largest outdegree in the causal network across diverse cell types (**Supplementary Figure 7a**), indicating the general importance of these factors in transcription regulation. We also experimented with the GET model trained using both quantitative ATAC signal and motif binding score as input and found similar top out-degree transcription factors (**Supplementary Figure 7b**).

Here we present four subnetworks in **Figure 4g** as examples. Notably, MBD2 and MECP2 show negative interaction with a promoter-enriched motif, GC-tract, which aligns with the well-known repressive effect of promoter methylation on gene expression^49,50^. The other three networks centered around NR/17 (representative transcription factor: ESR1), NFKB/1 (representative transcription factor: RELA), and ZFX exemplify the diverse information GET has learned. For example, the pair NR/17-Ebox/CACCTG highlights a functional regulatory complex ESR1-ZEB1^52^. NR/17-GLI is also supported by the known physical interaction between ESR1 and GLI3^53^. NFKB/1-Ebox/CACCTG has a strong interaction with negative effect size, while their representative transcription factors, RELA and SNAI1, have been shown to be interacting using co-immunoprecipitation^54^. ZFX is positively linked to TFAP2/1, and has been shown to co-localize with TFAP2A using ChIP-seq^55,56^ (**Supplementary Figure 7c**). The link between NFKB/1 and NFKB/2 is expected as they are commonly annotated to NFKB factors, despite having different sequence content.

To quantitatively assess the overlap with currently known physical interactions between transcription factors, we compared the GET motif-motif interaction network using two different approaches, GET Causal and GET Correlation, with the STRING v11^54^ database (**Methods:Causal discovery of regulator interaction**). Our results show a precision (true positive rate) of 5.6% by random chance. However, by selecting the top 1% (793 pairs) of causal or correlation pairs from GET’s predictions, we achieved precisions of 25.2% and 15.9%, respectively (**Figure 4h**). This confirms the advantage of our causal discovery based model interpretation approach. As a comparison, a recent mass spectrometry-based transcription factor-transcription factor interaction study^57^ reaches 30.4% precision with top 1.25% (990) pairs, while a colocalization score computed from HepG2 ChIP-seq for 677 transcription factors yields a larger number of motif-motif interactions yet has slightly lower precision and macro F1 score than GET Causal. Furthermore, GET Causal outperforms two accessibility-aware motif colocalization baselines, computed either across cell types or in hepatocytes. This reflects the incompleteness of annotated transcription factor-transcription factor interactions and highlights the value of GET-predicted motif pairs for identifying new transcription factor-transcription factor interactions.

### A structural catalog of human transcription factors and coactivators

With the predicted causal motif interaction network predicted by GET, we next embarked on building a structural catalog of the human transcription factor interactome using AlphaFold 2^58^. We started by categorizing transcription factor-transcription factor interactions into several different catalogs: Direct interactions, which includes homodimer, intra-family heterodimer, or inter-family heterodimer; and Cofactor-mediated interactions, which may encompass both cooperative and competitive binding (**Figure 5a**). Starting from the most straightforward intra-family interactions, we first acquired all dimeric structure predictions of more than 1,700 known human transcription factors. To evaluate whether AlphaFold predictions reflect true interactions, we assessed the result on predicting whether a transcription factor family can act as an “intra-family binder” based on the heuristic that intra-family binders should have a higher chance to form homodimers due to very similar structured domains. We found that AlphaFold can reach an area under the receiver operating characteristic (AUROC) of 0.69 and an average precision (AUPR) of 0.41 (**Supplementary Figure 8a**). The accuracy of AlphaFold dimer prediction is exemplified by the perfectly aligned TFAP2A structure to experimental results^59^ (**Supplementary Figure 8b**), even though there is no other similar template in PDB.

**Figure 5.**
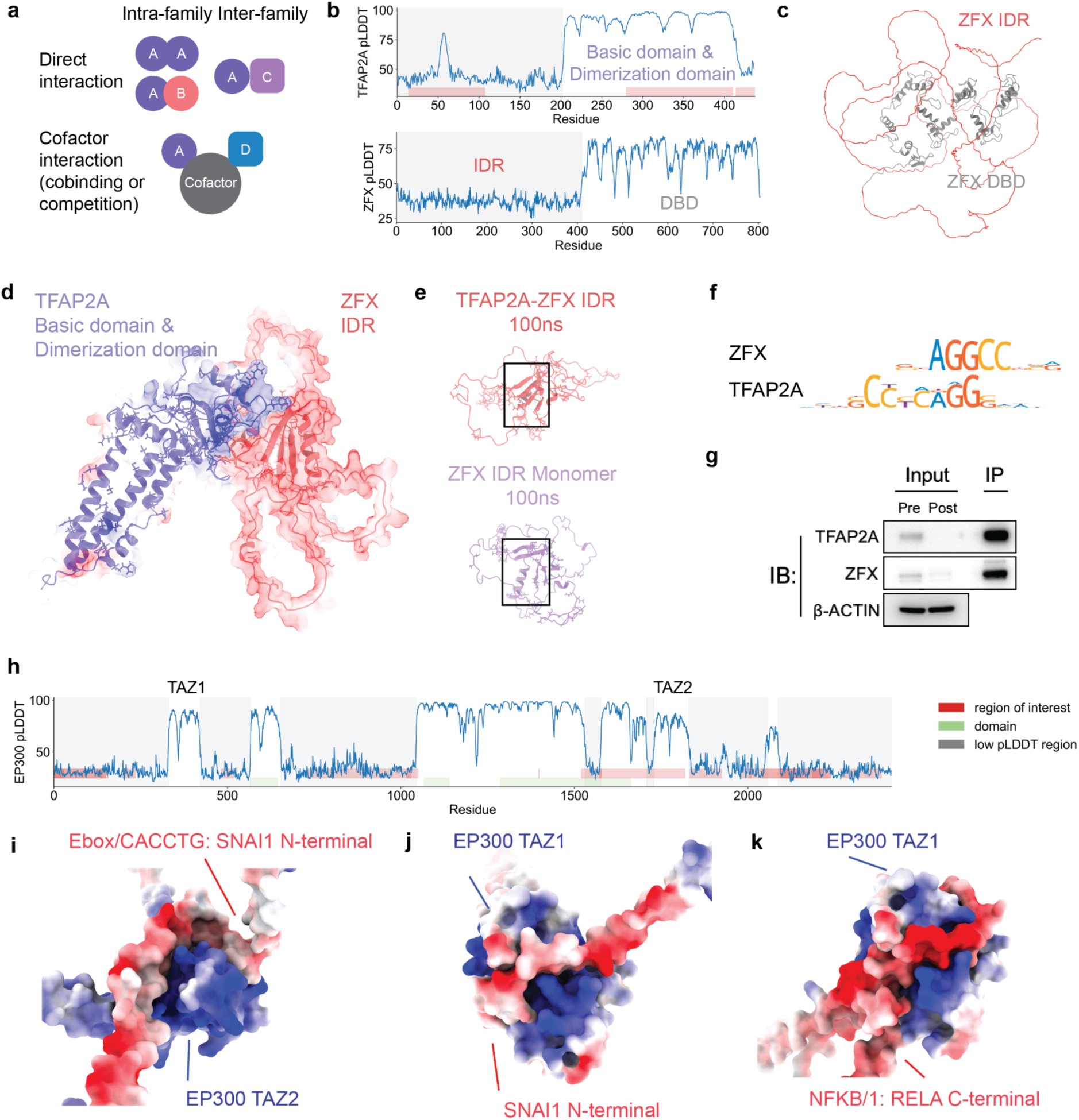
Structural properties of inferred transcription factor-transcription factor interactions through GET causal discovery. **a**. Catalogs of transcription factor-transcription factor interactions. Direct interactions include homodimer, intra-family heterodimer, or inter-family heterodimer. Cofactor-mediated interaction may include both cooperative and competitive binding. **b.** pLDDT plot for TFAP2A and ZFX, showing correspondence between high pLDDT regions and known protein domains (red rectangles). **c.** Predicted monomer structure of ZFX, showing DNA binding domain (DBD, grey) and intrinsically disordered region (IDR, red). **d.** Predicted structure of TFAP2A structured domains and ZFX IDR. Red and blue color marks negative and positive electrostatic surfaces. **e.** Molecular dynamics simulation of TFAP2A-ZFX IDR (red) or ZFX IDR monomer (purple). Collapsed structure in ZFX IDR monomer is highlighted in rectangle. **f.** Sequence logo of ZFX and TFAP2A transcription factor binding motifs. **g.** Co-immunoprecipitation analysis of TFAP2A and ZFX. **h**. pLDDT plot for EP300, highlighting TAZ1 and TAZ2 transcription factor interacting domains. Region of interest (red) and domain (green) marks annotated regions on UNIPROT. Low pLDDT regions are highlighted in gray shades. **i**. Prediction of structural interactions between SNAI1 N-terminal and EP300 TAZ2 domain. **j**. SNAI1 N-terminal and EP300 TAZ1 domain. **k**. RELA C-terminal and EP300 TAZ1 domain (right). Red and blue color marks negative and positive electrostatic surfaces.

With AlphaFold 2’s established ability to predict unseen multimer structures, we questioned whether the disordered region in the structure could fold upon binding to partners. Based on causal discovery predicted transcription factor-transcription factor interactions, we sought to identify potential structural interactions using AlphaFold 2. Taking TFAP2A and ZFX as an example, we segmented both proteins into four distinct structured or disordered domains based on predicted local distance difference test (pLDDT) (**Figure 5b**) and predicted the multimer structure of all pairwise combinations between these segments. Remarkably, the originally unstructured ZFX intrinsically disordered region (IDR) (**Figure 5c**) folded into a well-defined multimeric structure when paired with TFAP2A structured domains, mainly driven by electrostatic interactions (**Figure 5d**).

To provide another line of evidence, we employed molecular dynamics simulations (**Methods:Molecular dynamics simulation**), discovering that the monomer IDR exhibited a more collapsed structure after 100 ns (**Figure 5e**) and fewer inter-chain hydrogen bonds (**Supplementary Figure 8c**). Moreover, the per-residue pLDDT of ZFX, IDR, and TFAP2A (**Figure 5f**) in the multimer structure correlated strongly with residue instability, as measured by root mean squared distance (RMSD; **Supplementary Figure 8d**), aligning with previous studies indicating AlphaFold’s implicit learning of protein folding energy functions. To validate the predicted interactions between these two proteins, we performed co-immunoprecipitation experiments. As shown in **Figure 5g**, we were able to pull down ZFX using a TFAP2A antibody (**Methods:TFAP2A co-immunoprecipitation**).

When extending our method to negative effect pairs such as SNAI1 (Ebox/CACCTG) and RELA (NFKB/1), the absence of robust structural interactions led us to explore cofactor-mediated interactions, despite previous physical evidence (**Figure 5a, bottom**). Both transcription factors are known to physically interact with EP300^54^, and the predicted structures underscored electrostatic interactions with EP300’s TAZ1 and TAZ2 domains (**Figure 5h–k**). This concurs with existing studies on the electrostatic binding of transcription factor IDR to EP300 TAZ domains^60–63^.

Broadening our study, we applied the procedure to top 5% transcription factor pairs in each cell type (totaling 1,718 transcription factor pairs or 24,737 pairs of transcription factor segments, see **Methods:Causal discovery of regulator interaction**) as predicted through GET-based causal discovery and built a structural catalog of transcription factor interactions. Interestingly, the folded conformation of ZFX IDR can also be seen in other transcription factor pairs, for example EGR1 IDR-ZFX IDR (**Supplementary Figure 8e**). We also show that the previously mentioned interaction between ESR1 and ZEB1 could be driven by a confident structural interaction between the ZEB1 C-terminal IDR and ESR1 NR domain (**Supplementary Figure 8f**).

### Exploring the mechanism of a leukemia-predisposing germline mutation in PAX5 IDR with GET

To demonstrate the utility of information provided by the GET Catalog, we performed a case study on PAX5, a driver transcription factor of B-cell precursor acute lymphoblastic leukemia (B-ALL)^64^. B-ALL is the most frequent pediatric malignancy, and somatic genetic alterations (deletions, translocations, and mutations) in PAX5 occur in approximately 30% of sporadic cases^65^. While most PAX5 somatic missense mutations affect the DNA-binding domain (V26G or P80R), G183S is a recurrent familial germline mutation that confers an elevated risk of developing B-ALL^64–66^. Somatic mutation of G183 and frameshift in a nearby hotspot are also seen in B-ALL patients^67^. Although the pLDDT plot of PAX5 highlights G183 and the octapeptide domain as a small peak in the entire intrinsically disordered region, its functional role remains elusive (**Figure 6a**).

**Figure 6.**
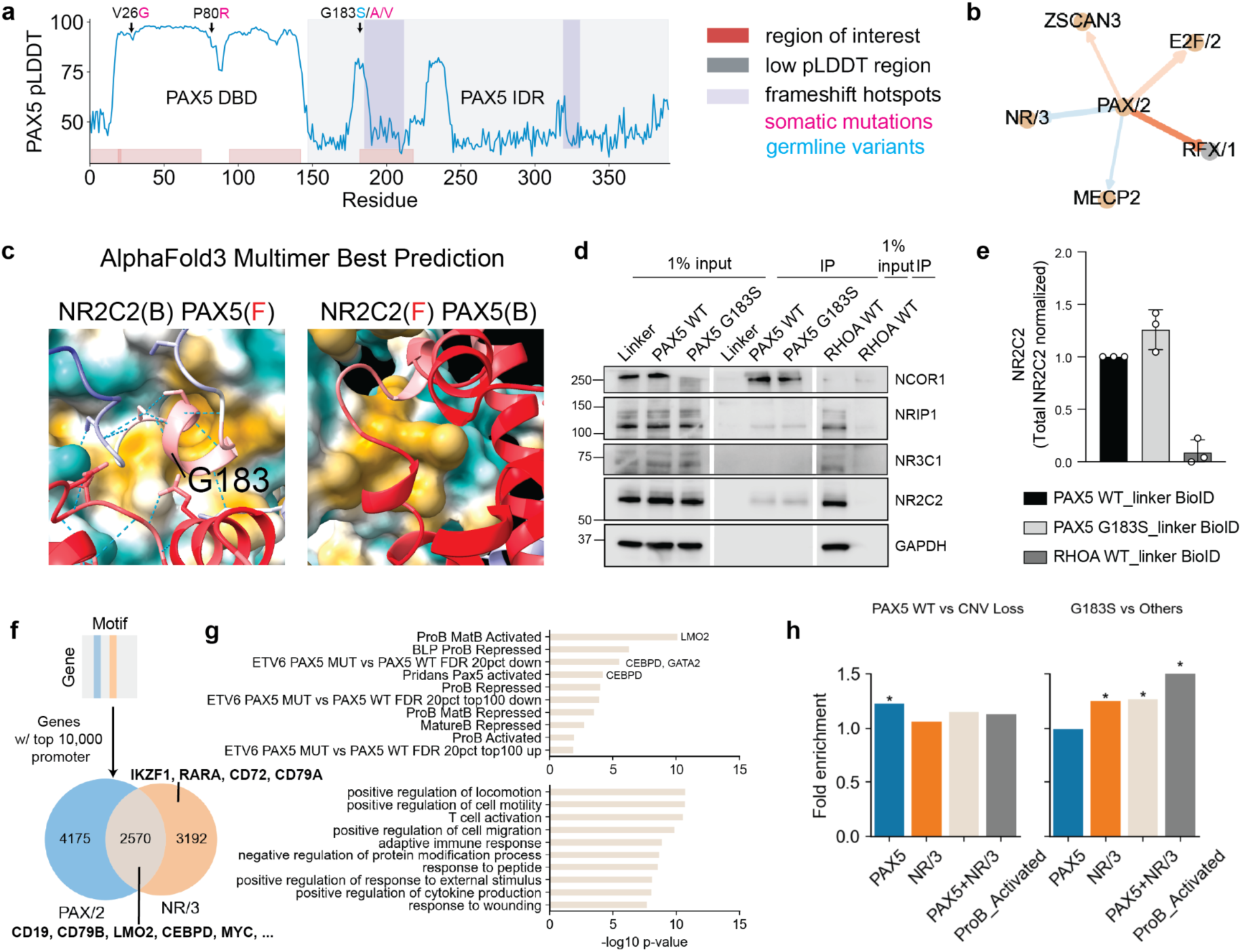
GET identifies a cell type specific transcription factor-transcription factor interaction affected by a cancer-associated germline variant. **a**. pLDDT plot for PAX5. Showing three mutational hotspots: V26G, P80R, G183S/V/A, and two frameshift hotspots^67^. Region annotations from UNIPROT are shown in the figure as “region of interest.” **b**. B-cell specific motif interactions of PAX/2. PAX5 is the highest expressed transcription factor with PAX/2 motif. RORA is the highest expressed transcription factor with the NR/3 motif. Color scheme follows Figure 4g. **c.** AlphaFold 3 predicted multimer structure of PAX5 IDR and RORA NR domain showing contacts around G183 (B: Back; F: Front). The blue-yellow surface in the back shows hydrophilicity and hydrophobicity, respectively. Blue-red strands in the front show low-high prediction confidence, respectively. **d.** Detection of NCOR1, NRIP1, NR3C1, and NR2C2 PAX5 interacting proteins in immunoprecipitates from PAX5 WT, PAX5 G183S and RHOA WT-BioID-expressing B-ALL REH cell line and in total protein lysates using Protein Ligation Assays. A representative experiment is shown. **e.** Quantification of PAX5-NR2C2 interaction in the streptavidin immunoprecipitation shown in d. **f.** Venn diagram of identified PAX/2 and NR/3 specific and common regulatory targets using GET gene-by-motif importance matrix. **g.** (Top) Enrichment analysis (-log 10 p-value from Fisher exact test) using B-cell associated gene sets in Shah et al.^64^ and (bottom) biological process gene ontology gene sets. Results for the PAX/2-NR/3 common genes are shown in this figure. Results for PAX/2 or NR/3 specific genes are shown in **Supplementary Figure 9**. **h.** Enrichment analysis for differentially expressed genes between PAX5 wild type vs. PAX5 loss (left) and PAX5 G183S vs. other PAX5 alterations (CNV loss, P80R). * indicates statistical significance from hypergeometric tests. Benjamini-Hochberg adjusted P-values are reported.

To probe this, we first explored potential interaction pairs involving PAX5 (PAX/2 motif) in fetal B lymphocytes (CXCR5+). We identified promising interactions with several transcription factors including E2F3, MZF1, MECP2, NR4A2, RFX3, and RORA (NR/3 motif, **Figure 6b**). Subsequent exhaustive segment interaction screening revealed a novel interaction between the RORA nuclear receptor (NR) domain and the octapeptide domain of PAX5, confirmed by both AlphaFold 2 and AlphaFold 3 (**Figure 6c**). The binding interface of PAX5 IDR and NR domain show hydrophobicity on both sides. The G183 residue is close to the binding site and a series of Serine residues linked by hydrogen bonds. Thus, the Glycine to Serine mutation could potentially affect the interaction with the NR domain leading to changes in the coregulated transcriptional program. This interaction was further corroborated by positive affinity purification-mass spectrometry (AP-MS) data of their paralog PAX2-NR2C2^57^, as both the PAX5 octapeptide domain and NR domain are highly conserved and structurally similar across their paralogs.

To validate the interaction between PAX5 and NR/3 motif-containing proteins, we selected candidate NR/3 transcription factors by prioritizing all TFs in the NR/3 motif family based on their expression in 15 B-ALL PAX wildtype patient samples in a longitudinal study. We found the highest expressed NR/3 transcription factor to be NR4A1 and NR2C2, with NR2C2 showing less variable expression across patients. We performed Protein Ligation Assays using PAX5 wild type and G183S mutant linked to a BioID biotin ligase in the B-ALL REH cell line. Analysis of proteins biotinylated by their proximity to the PAX5-BioID fusions identified the nuclear corepressor NCOR1, a previously described PAX5 interactor^68,69^, NR2C2 NR/3 motif-containing nuclear corepressor, and the NRIP1 nuclear receptor interacting protein. The experiment shows a clear interaction between PAX5 and NR2C2, aligned with the previous report, while NR4A1 shows no interaction with PAX5. NR3C1, a NR/20 motif TF predicted to be interacting with NR/3 but not PAX/1 by GET, shows no interaction with PAX5. Interestingly, the presence of the G183S mutation results in an increased interaction between PAX5 and NR2C2 (**Figure 6d-e**).

To elucidate whether the PAX/2 and NR/3 motifs coregulate genes, we examined the top 10,000 promoters predicted to be most influenced by them. Our analysis uncovered a set of 2,570 genes commonly regulated by both, including surface markers like CD19 and CD79B, as well as known oncogenes implicated in B-ALL including MYC, CEBPD, and LMO2, although these oncogenes are also predicted to be strongly repressed by IKZF1 (IKAROS tumor suppressor, with ZNF143 motif) and are not highly expressed (**Figure 6f-g**). Enrichment analysis revealed an overrepresentation of genes involved in leukocyte activation and genes affected by PAX5 perturbation during B cell differentiation, aligning with previous work on the G183S mutation^64,70–75^ (**Figure 6e, Supplementary Figure 9a-b**). On the other hand, the genes that are specifically regulated by PAX/2 or NR/3 are enriched in neuronal pathways and the cell cycle, respectively.

To investigate the effect of the PAX5 G183S mutation on transcriptional programs of NR/3 and PAX5 regulated genes, we examined 141 sporadic childhood B-ALL samples, including 4 familial B-ALL samples with PAX5 G183S germline variants. We first selected only CDKN2A loss cases (n=20) to match with familial cases (n=4, biallelic mutations, loss of WT PAX5 and CDKN2A). We further stratified the patients into two groups to perform two different comparisons: 1) PAX5 WT (n=10) vs. PAX5 Loss (n=10), as a negative control; and 2) PAX5 G183S (n=4) vs. PAX5 loss of function cases (e.g. PAX5 P80R, PAX5 Loss, n=12) for defining the G183S-specific transcriptome signature. The analysis identifies a specific transcriptional program associated with the G183S mutation (**Supplementary Figure 9e**). To evaluate if this specific transcriptional program was relevant to PAX5-NR interaction, we performed further gene set enrichment analysis, finding an enrichment in the PAX5-loss associated differential expression genes (fold enrichment 1.22, P-value=1.5e-3, hypergeometric test with Benjamini-Hochberg correction) with predicted PAX5 targets and no significant enrichment in NR/3 regulated and PAX5+NR/3 regulated genes, as expected. However, the PAX5 G183S specific differential expression genes show significant enrichment in NR/3 and PAX5+NR/3 regulated genes (fold enrichment: 1.41, 1.35; P-value=1.05e-30, 1.57e-18). Moreover, we found that this mutant specific differential expression gene set is also enriched in the genes activated in ProB cells (fold enrichment: 1.57, P-value=2.2e-3), including well-known functionally relevant genes like CD19, EBF1, BACH2, and PML (**Figure 6g**).

## Discussion

In this study, we introduced GET, a state-of-the-art foundation model specifically engineered to decipher mechanisms governing transcriptional regulation across a wide range of human cell types. By integrating chromatin accessibility data and genomic sequence information, GET achieves a level of predictive precision comparable to experimental replicates in leave-out cell types. Furthermore, GET demonstrates exceptional adaptability across an array of sequencing platforms and assay types, as well as non-physiological cell types like tumor cells. Through interpretation of the model, we successfully identified long range regulatory elements in fetal hemoglobin and their associated transcription factors. Collecting regulatory information from all 213 cell types and synergizing TF-TF interactions deduced through GET with protein structure predictions, we constructed the publicly accessible GET Catalog (https://huggingface.co/spaces/get-foundation/getdemo). Utilizing the PAX5 gene as a case study, we illustrated the catalog’s utility in elucidating functional variants in disordered protein domains that were previously difficult to study.

Current limitations of GET include a reliance primarily on chromatin accessibility data, bounded resolution to distinguish between transcription factor homologs that have very similar motifs, and training on only coarse-grained cell states and region-level sequence information. Future enhancements to GET can be envisioned through the incorporation of multiple layers of biological information, including but not limited to nucleotide-level regulator footprints^5,26,76^, three-dimensional chromatin architecture^77,78^, and regulator expression profiles or single cell embeddings^11–13^. Future iterations of GET can incorporate a broader range of assays, including those that directly measure transcription factor binding, histone modifications, and RNA polymerase activity, to provide a more holistic view of the regulatory landscape.

Multiplexed nucleotide-level perturbations or randomizations will be instrumental in calibrating GET for precise prediction of the functional impact of noncoding genetic variants. Elucidating the effect of non-coding variants in modulating gene expression and disease susceptibility remains an important area of exploration. Integrating genomic variants into the GET framework will enable prediction of their impact on gene regulation more accurately, offering insights into the genetic basis of complex traits and diseases. Additionally, the kinetics of gene regulation, reflecting the temporal changes in transcriptional activity in response to developmental cues or environmental stimuli, is another dimension of complexity that can be integrated into the model. As in our PAX5-NR2C2 example, other types of information like TF expression in specific cell types or protein-protein interaction data can help to further narrow down specific transcription factors within motif-motif interaction networks predicted by GET. With our efficient finetuning framework, comparative interpretation analysis using pretrained and finetuned GET can be performed to highlight important regulatory regions or motifs driving cell state changes.

Leveraging GET as a computational framework, generative models can be developed to design megabase-scale enhancer arrays and engineer cell-type specific transcription factors or their interaction inhibitors for targeted therapeutic interventions. Extending GET’s pretraining set to include data from cellular perturbations, such as drug treatments or genetic modifications, or from cancer cells, will enhance the model’s ability to predict the outcomes of experimental interventions. This may facilitate the design of therapeutic strategies and the identification of novel drug targets. Collectively, GET represents a pioneering approach in cell type-specific transcriptional modeling, with broad applicability in the identification of regulatory elements, upstream regulators, and crucial transcription factor interactions.

## Data availability

Bulk RNA-sequencing of B-ALL patients published in our previous study^68^ is acquired at SRA (PRJNA534488). Human transcription factor protein interaction networks are downloaded from supplementary data of Göös et al.^57^ Precomputed regulatory inference results, preprocessed data, and structure predictions can be viewed at the GET website: https://huggingface.co/spaces/get-foundation/GET. The full processed data and inference results are provided in a public AWS S3 bucket.

## Code availability

Code for pretraining, finetuning, data preprocessing, and analysis will be made available in GitHub (https://github.com/RabadanLab, https://github.com/GET-Foundation) after publication. The pretrained model will be available on Hugging Face (https://huggingface.co/get-foundation). Code for the website is available at https://huggingface.co/spaces/get-foundation/GET/tree/main.

## Author contributions

X.F. and S.M. initiated the project. X.F. and R.R. conceived of the study and designed the analyses. X.F. and S.M. designed the model with advice from E.X. X.F. and S.M. implemented the model. X.F. performed data processing. S.M., X.F. and A.B. performed model training, ablation and performance analyses. X.F. performed model interpretation analysis including MPRA, regulatory elements and regulator prediction, network analysis, and structural analysis. X.F. and S.M. constructed the GET catalog. X.F., S.M., and A.B. built the website. A.S. and A.L. performed the experimental validation with input from D.O. and T.P. M.M.A, J.S., R.S., and T.J. helped with analysis. X.F. and J.T. co-designed an updated version of the data processing pipeline. Y.L. provided suggestions and computational resources to a pilot study. A.C. provided critical suggestions to the analysis. A.F. provided critical suggestions and edited the manuscript. E.X. and R.R. supervised the study. X.F., S.M., A.B., M.M.A., E.X., and R.R. prepared the manuscript with input from all authors.

## Acknowledgements

We gratefully acknowledge funding from NIH (R35 CA253126 to RR, P01 CA174653 to RR, R01 HL159377 to A.F. and R.R., U01 CA243073 to R.R. and T.P.) and SU2C Convergence 3.14 to RR.

---

**Supplementary Figure 1.**
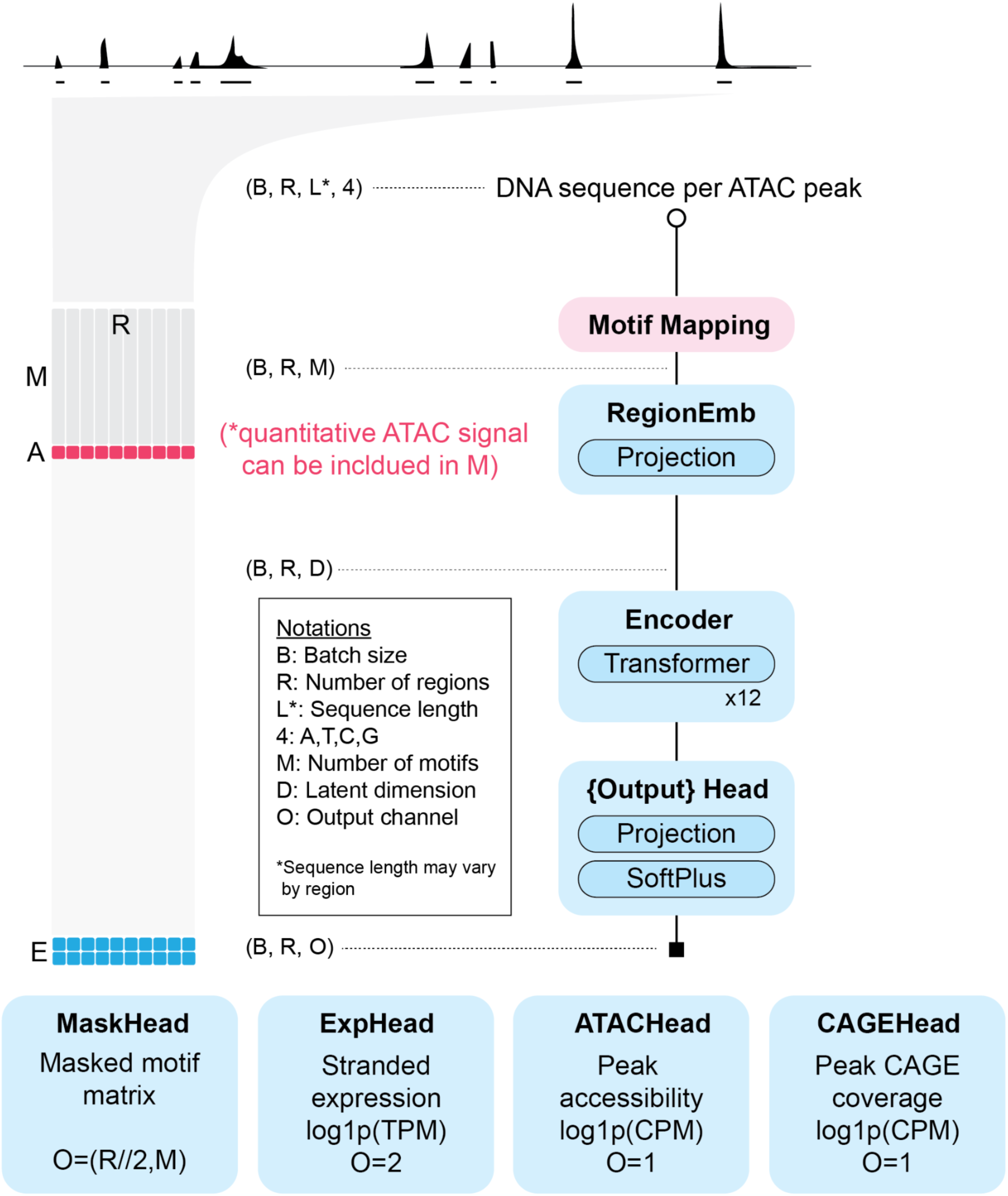
Architecture of GET.

**Supplementary Figure 2.**
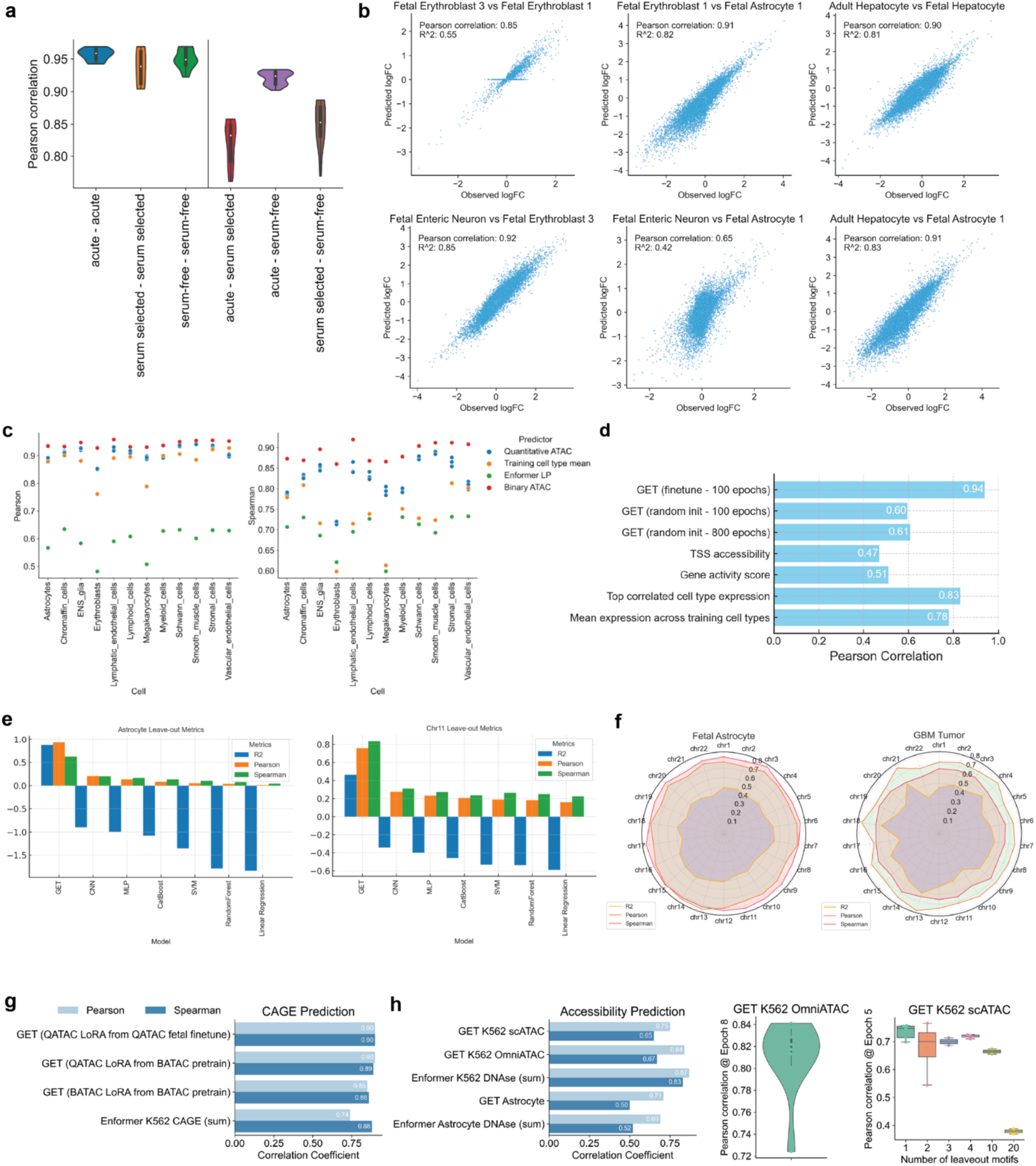
GET model performance and benchmarking. **a.** Pearson correlation of gene expression between biological replicates and across different culture systems for human astrocytes. **b.** Scatterplots showing predicted vs. observed log fold change (log FC) in gene expression between different cell type pairs. Top row compares erythroblast subtypes and hepatocytes. Bottom row compares enteric neurons to erythroblasts and astrocytes. Pearson correlations and R^2^ values are provided for each comparison. **c** Comparison of Pearson correlation between predicted and observed gene expression in left-out astrocytes for different models and baselines. <u>Quantitative ATAC</u>: GET model with quantitative ATAC and motif input. <u>Binarized ATAC</u>: GET model with only motif input. <u>Enformer LP</u>: Linear probing of Enformer CAGE header outputs. <u>Training cell type mean</u>: Mean expression value across all training cell types. GET with finetuning shows the highest correlation at 0.94. **d.** Ablation study of GET pre-training, showing leave-out-astrocyte finetuning performance. **e**. Comparison of GET to baseline machine learning models (CNN, MLP, CatBoost, SVMRegression, RandomForest, and LinearRegression) on leave-out-chromosome 11 and leave-out-astrocyte prediction performance (R^2^, Pearson correlation, Spearman correlation). **f.** Radar plot showing leave-one-chromosome-out finetuning performance (R^2^, Pearson correlation, Spearman correlation) of GET in Fetal Astrocyte and GBM tumor samples. **g**. Three settings of GET finetuned on CAGE K562 dataset and evaluated on leave-out chromosome 14 as compared to Enformer predictions. GET finetuned with quantitative ATAC (“QATAC LoRA from QATAC fetal finetune” and “QATAC LoRA from BATAC pretrain”) outperforms GET finetuned with binarized ATAC (“BATAC LoRA from BATAC pretrain”) and Enformer predictions for the peak evaluation regions. Evaluation was performed for all TSS in chromosome 14. **h**. Performance of GET on ATAC prediction. (left) GET performance for K562 and Astrocyte ATAC compared against Enformer’s predictions for DNAse on K562 and Astrocyte output tracks. (middle) GET performance for K562 bulk OmniATAC, showing leave-one-chromosome-out results for all autosomes. (right) GET performance for K562 scATAC, showing leave-out-motif analysis for randomly selected 1, 2, 3, 4, 10, or 20 motif features from the training set and evaluating on peaks with the left-out motifs (**Method: Model evaluation**).

**Supplementary Figure 3.**
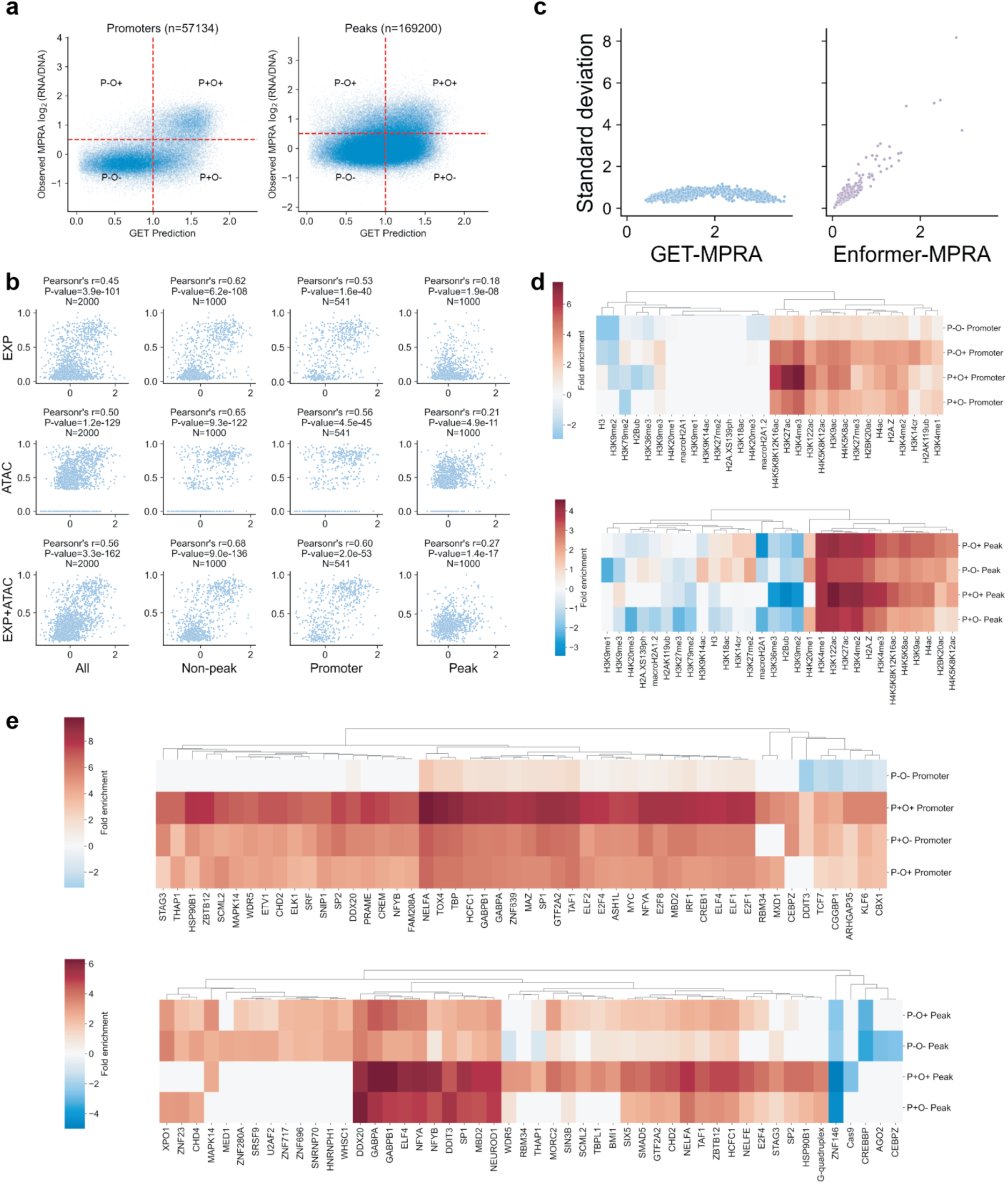
**a**. Scatter plot of lentiMPRA readout and GET-MPRA prediction. Promoter (left) or ATAC peak (right) elements are gated into four sub-categories, respectively, based on high (+) or low (-) in <u>P</u>rediction (cutoff=1) or <u>O</u>bservation (cutoff=0.5). **b**. Detailed ablation of expression and ATAC components of GET prediction. **c**. Benchmarking standard deviation of MPRA zero-shot prediction across random insertions. **d.** Histone mark enrichment analysis of promoter (Top) and peak (Bottom) elements respectively using ENCODE K562 ChIP-seq data. **e.** Transcription factor binding site enrichment analysis of promoter (Top) and peak (Bottom) elements respectively using ENCODE K562 ChIP-seq data. Fisher exact test was performed. Tests with a p-value<0.05 are shown. Color shows log10 (Fold enrichment). For transcription factors, the variance of fold enrichment cross four groups was calculated, and the top 50 TFs are visualized.

**Supplementary Figure 4.**
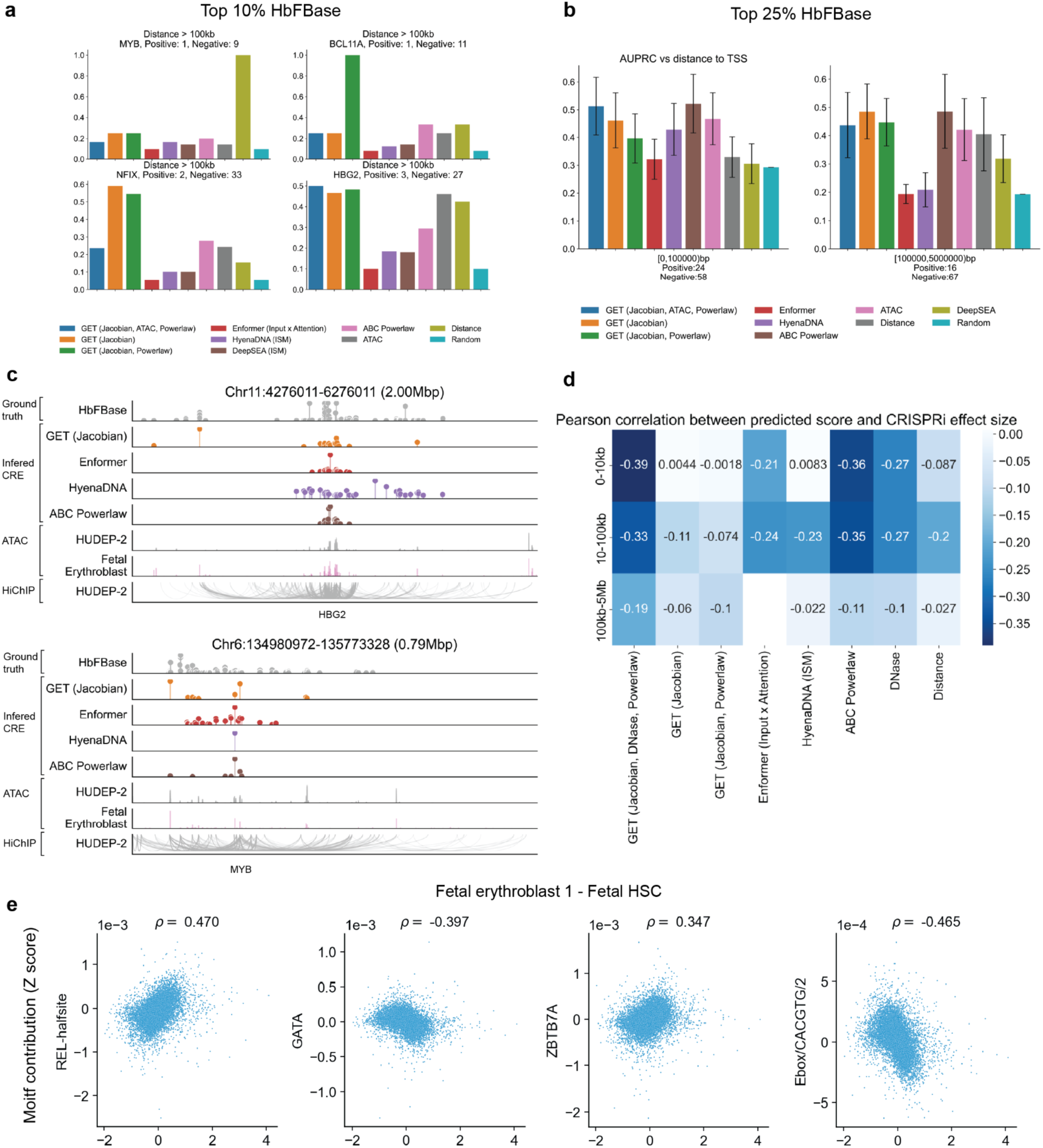
**a.** Per-gene AUPRC benchmark of cis-regulatory region prediction with top 10% HbFbase as the label cut off. **b.** Distance stratified AUPRC benchmark using top 25% HbFbase as the label cut off. **c.** Cis-regulatory region prediction of HBG2 and MYB loci. **d.** Distance-stratified Pearson correlation between predicted scores and K562 CRISPRi effect size. **e**. Correlation between motif contribution (y-axis) in Fetal Erythroblast 1 and the predicted target gene expression change (x-axis) between Fetal Erythroblast 1 and Fetal HSC. Four motifs relevant to erythroid differentiation are shown.

**Supplementary Figure 5.**
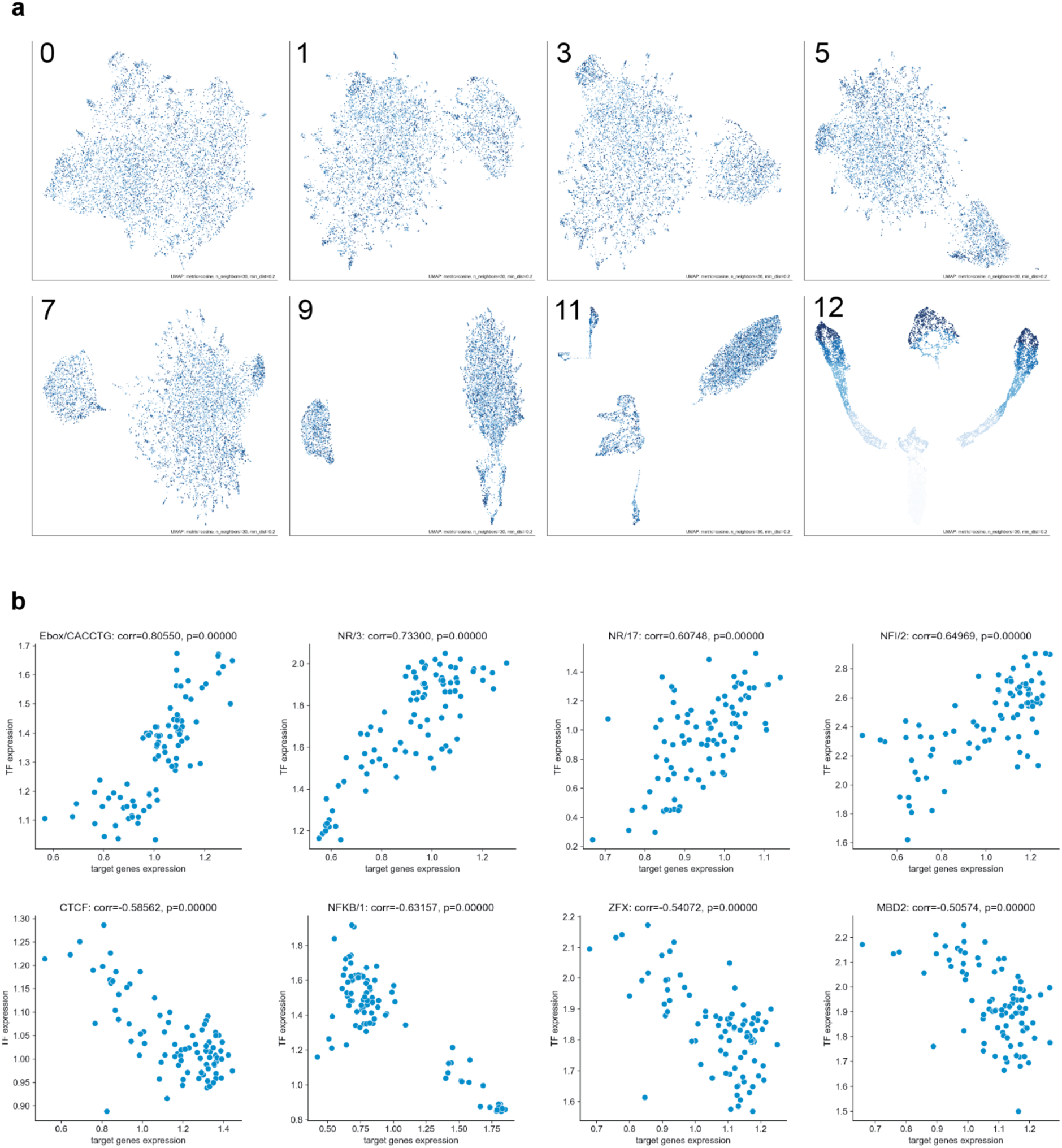
**a.** Example latent embedding space after 12 transformer layers. Each dot is a gene colored by expression value. **b.** Correlation between mean transcription factor expression in a motif cluster versus the mean expression of their target genes predicted by GET across fetal cell types.

**Supplementary Figure 6.**
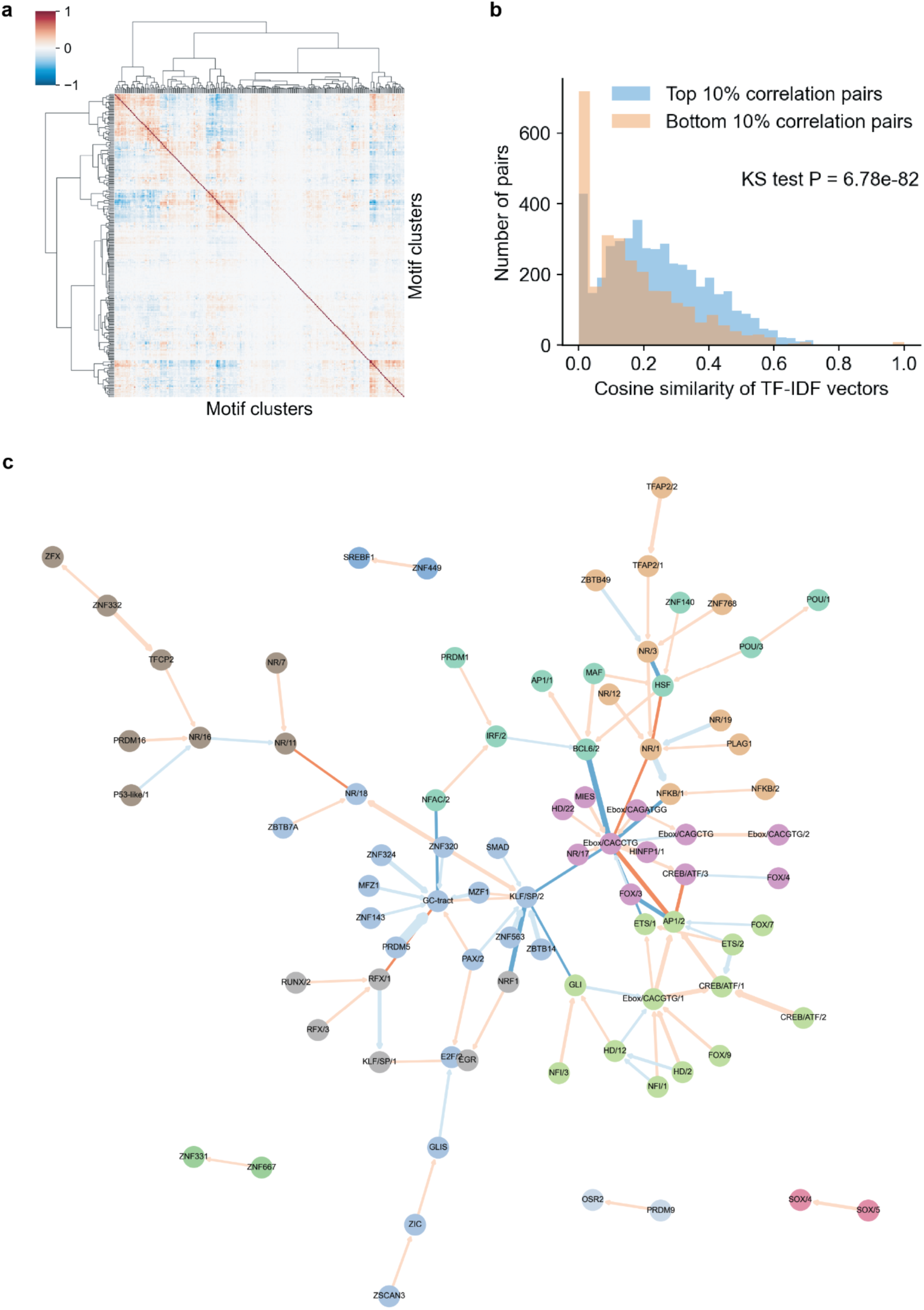
**a.** Correlations between motif clusters using GET’s Jacobian matrix show potential co-regulating motifs. **b.** Top correlated motif pairs have significantly larger functional similarity. X-axis is cosine similarity computed on term (motif clusters) frequency– inverse document (Gene Ontology biological process) frequency (transcription factor-IDF) matrix. **c**. Casual motif-motif interaction network inferred from all cell types. Edges with absolute effect size smaller than 0.1 and isolated nodes are removed for visualization purposes. Node colors represent communities. Edge weights represent causal effect sizes. Edge colors represent positive (orange) or negative (blue) effects.

**Supplementary Figure 7.**
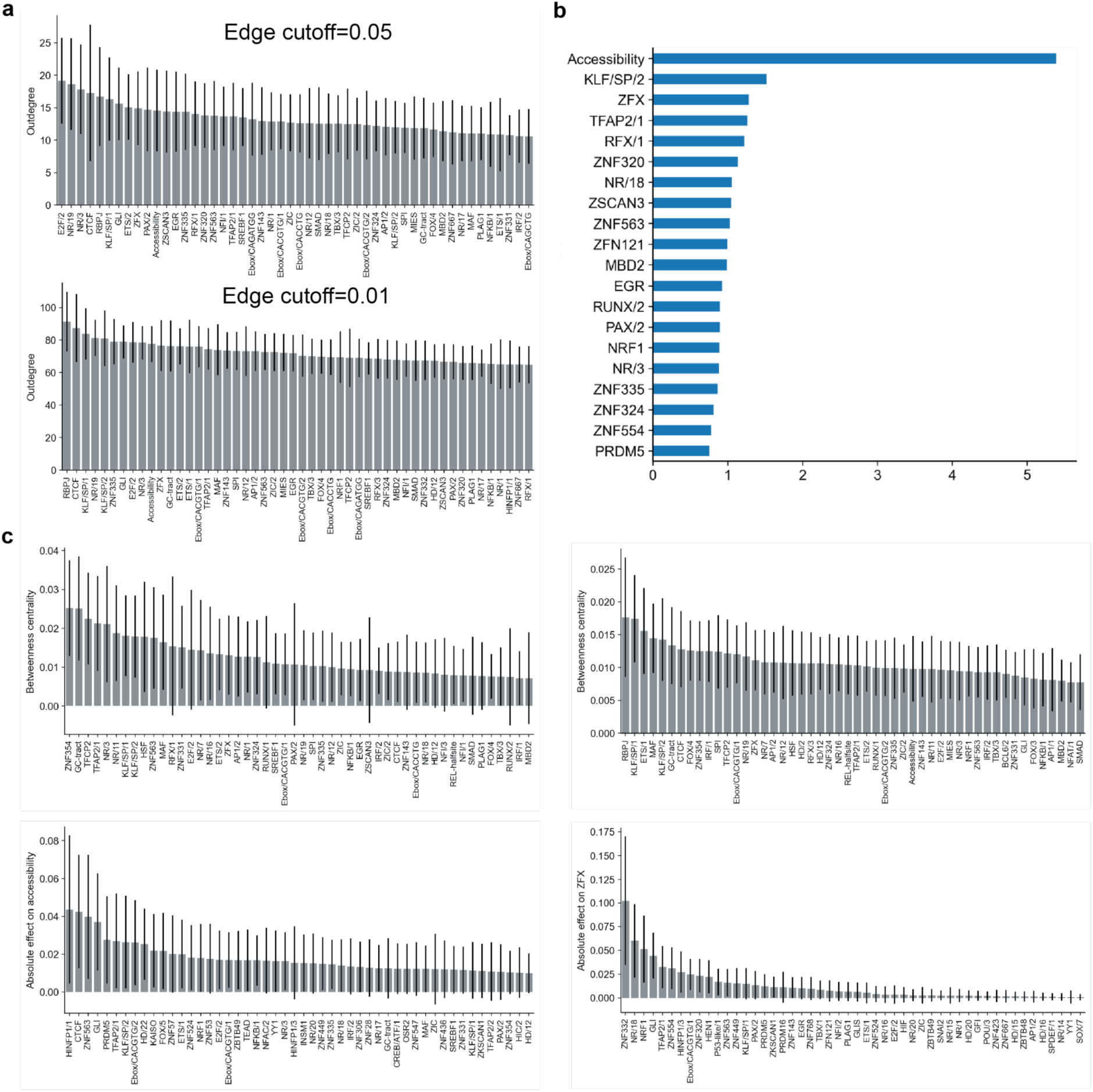
Overall network statistics of causal motif-motif network. **a.** Out-degree distribution across cell-type-specific causal networks at two different absolute edge weight cutoffs. Error bar shows standard deviation across cell types. **b.** Out-degree distribution of Quantitative-ATAC GET model trained using fetal data. **c.** Betweenness centrality and absolute effect on accessibility or ZFX across cell-type-specific causal networks at edge cutoff 0.01.

**Supplementary Figure 8.**
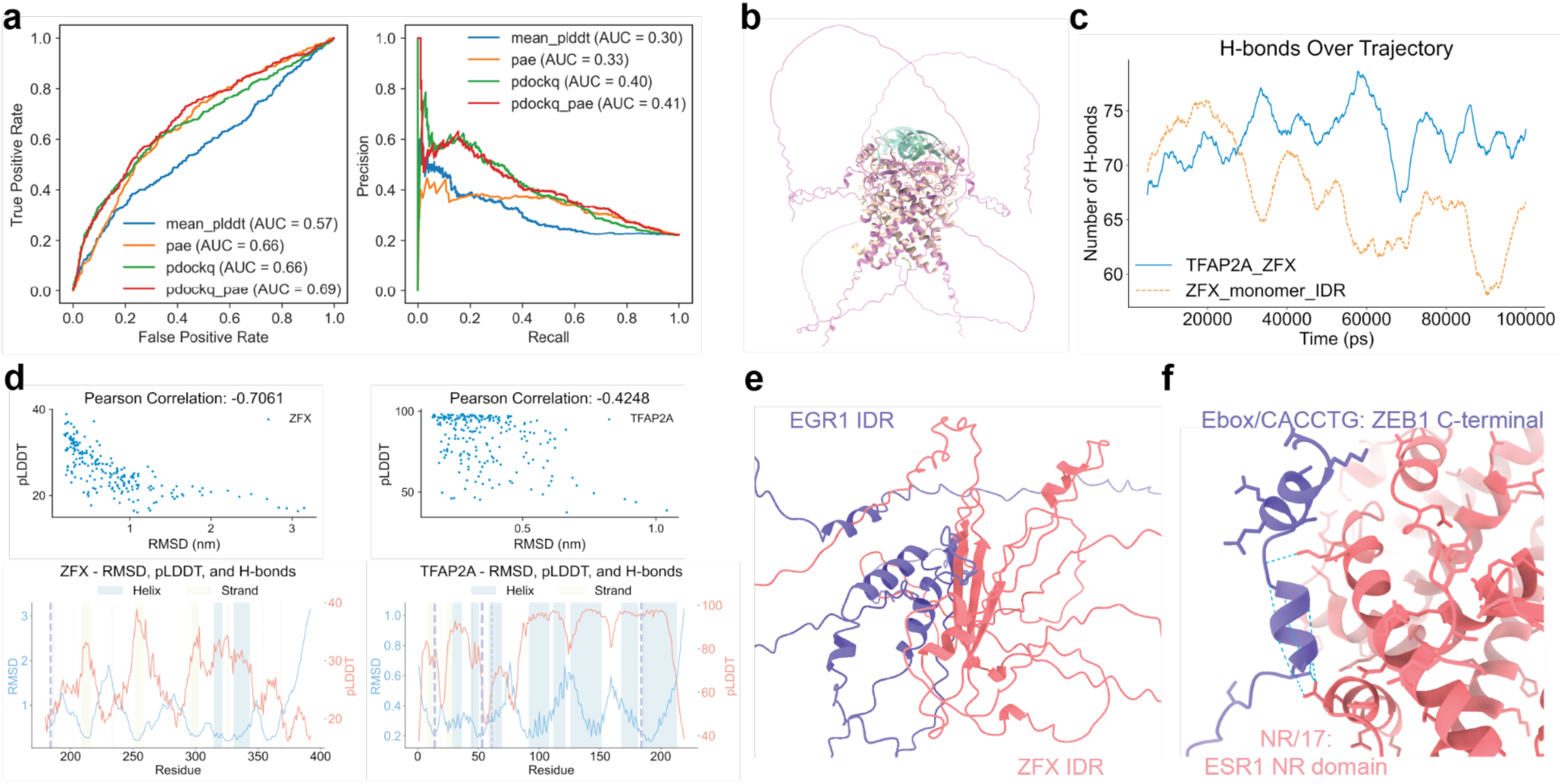
**a**. Prediction performance of “intra-family binder” using different multimer structure prediction confidence scores. (Left) ROC curve with x-axis showing false positive rate and y-axis showing true positive rate. (Right) PR curve with x-axis showing recall and y-axis showing precision. <u>mean_plddt</u>: Average predicted Local Distance Difference Test (pLDDT) score across all residues. <u>pae</u>: Predicted Aligned Error across all inter-chain interactions. <u>pdockq</u>: Predicted DockQ metric using interface pLDDT. <u>pdockq_pae</u>: Multiplication of pDockQ and pAE. **b**. Comparison of AlphaFold 2 predicted TFAP2A dimer structure (pink) with crystal structure of TFAP2A (yellow)-DNA (green) complex. **c.** Change in number of hydrogen bonds in TFAP2A-ZFX IDR complex or ZFX IDR monomer across simulation trajectory. **d.** Correlation between pLDDT and residue RMSD across the simulation trajectory of ZFX IDR in the complex structure. Visualized in scatter plot (Top) and line plot across the protein sequence (Bottom). Yellow and blue shades in the line plot highlight beta sheets or alpha helices. **e**. Predicted multimer structure of EGR1 IDR-ZFX IDR. **f.** Predicted multimer structure of ZEB1 C-terminal and ESR1 NR domain.

**Supplementary Figure 9.**
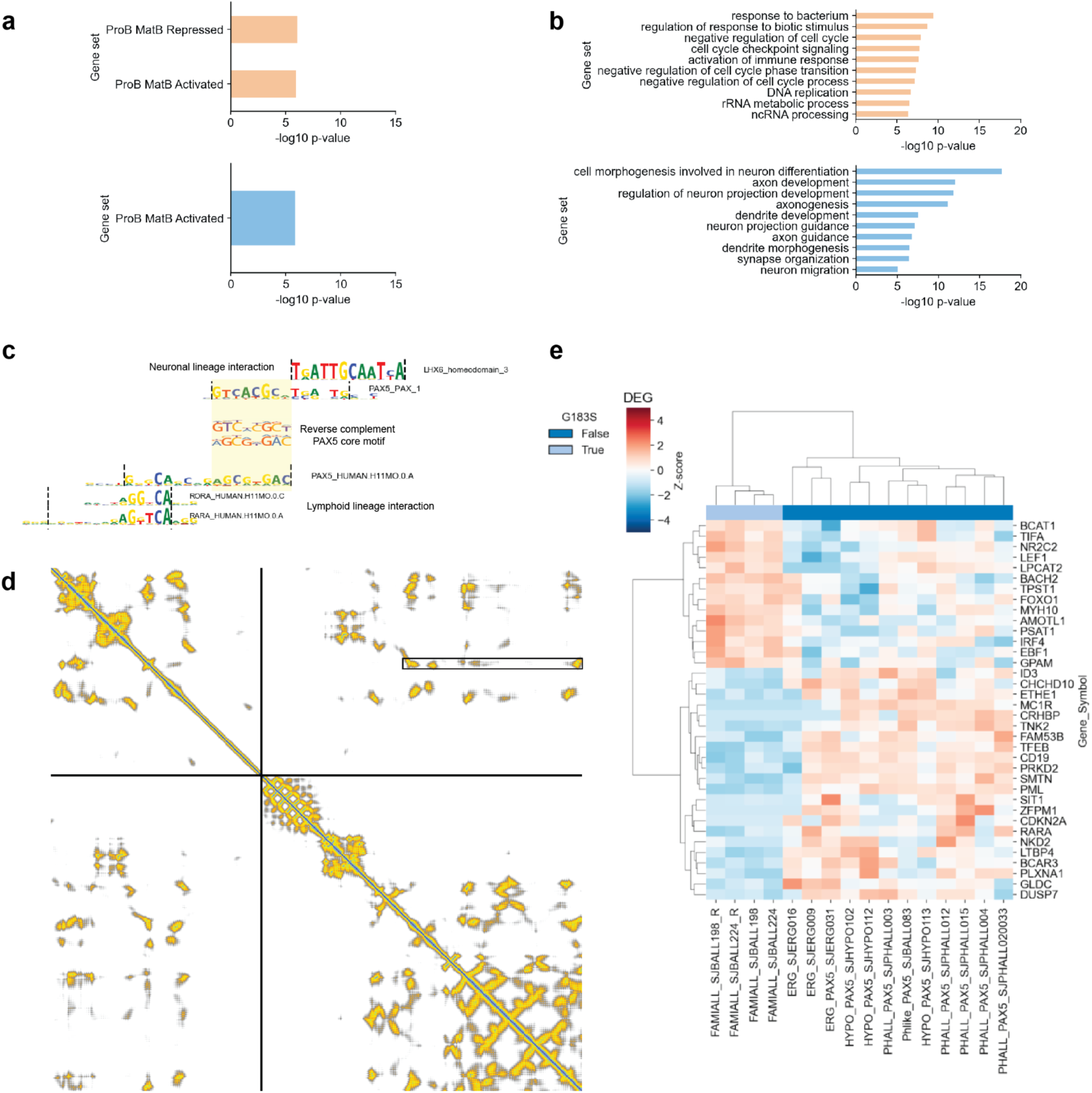
**a.** Gene enrichment analysis of NR/3-specific (orange) and PAX/2-specific (blue) target genes using published B cell related gene sets^64^ and **b.** gene ontology biological processes. **c.** Motif visualization and comparison. From top to bottom: LHX6 (a neuronal lineage TF), PAX5 motif in PAX/1 motif cluster, PAX5 core motif and reverse complement, PAX5 motif in PAX/2 motif cluster, RORA and RARA motifs in NR/3 motif cluster. **d.** Contact map of full length PAX5-NR2C2 from AlphaFold 3 prediction. The best ranked model is shown here. G183 residue is highlighted in the black box. **e.** Heatmap with the specific transcriptional program for the cases with the germline PAX5 G183S mutation.

