## Supplementary material for "GET: a foundation model of transcription across human cell types": Method

### Methods

#### ATAC-seq data processing

##### Pseudobulk

To determine the chromatin accessibility score for each region, we utilized the scATAC-seq count table and the cell annotation table from each respective study. To aggregate single cells into "pseudobulk cell types," we used the Louvain clustering results from each study. Log count per million was used for each pseudobulk. The annotations of each cluster were used to determine the biological cell type. Empirically, we set a threshold of more than 600 cell counts to ensure adequate sequencing depth for selected cell clusters. A comprehensive table of pseudobulk cell types used during the training process can be found in Supplementary Table S1.

In summary, we used ATAC-seq and expression data from references[1–3]. In total, the dataset encompasses 1.3 million single nuclei. The data is only presented in pseudobulked format. All cell types are primary cell types from normal tissue. No disease states were included in the pretraining dataset. We incorporated additional datasets in downstream tasks like K562 and zero-shot analysis in hESC and tumor cells.

##### Cell-type-specific accessible region identification

For the identification of cell-type-specific accessible regions, the peak calling results from the original studies of each dataset were followed to obtain a union set of peaks. Subsequently, to compile a list of accessible regions specific to each cell type, peaks with no counts were filtered out.

In the context of the human fetal and adult chromatin accessibility atlas, we used the peak set produced by Zhang et al.[2], incorporating the fetal chromatin accessibility atlas originally published by Domcke et al.[4]. We have also trained a version of the fetal-only GET model using the original peak calling and cell type annotation from Domcke et al.[4], resulting in comparable expression prediction and regulatory analysis performance. For the 10x multiome data, we used the provided peak fragment count matrix. For the K562 NEAT-seq and bulk chromatin accessibility data, a more permissive version of peaks was called using MACS2[5], and different logTPM cutoffs were applied to the resulting peak set to select accessible regions. This accessibility-based data augmentation enhances the diversity of input data and finetunes the GET model for data from a single cell type.

##### Accessibility features

In our study, the chromatin accessibility score for a specific genomic region is defined by the count of fragments located within that region for a given cell type pseudobulk. To enhance the model's generalizability, these counts are further normalized through the logCPM (Log Counts Per Million) procedure. Specifically, let $t$ be the total fragment count in a pseudobulk, and $c_i$ be the fragment count in region $i$. Then, the accessibility score $s_i$ is computed as:

$$s_i = \log_{10}\left(\frac{c_i}{t}+1\right), \quad t = \sum_i c_i$$

For the majority of the regulatory analysis, the "Binary-ATAC" version of the GET model is utilized to comprehensively evaluate the regulatory influences exerted by transcription factors. In both the training and inference phases of this specific model version, the accessibility scores are uniformly set to 1 if the region is identified as a chromatin accessibility peak. This equates to assuming binary chromatin accessibility states within the studied scenario.

##### Motif features

To calculate the motif binding score within a specific genomic region, the corresponding sequence is scanned against the hg38 reference genome. This procedure involves utilizing 2,179 transcription factor motif position weight matrices (PWMs), as previously compiled by Vierstra et al.[6], accessible at Vierstra's resources. For the scanning process, the MOODS tool is used with default threshold[7].

More specifically, to represent sequence information while mitigating feature redundancy, a specialized motif scoring process is implemented. Building on Vierstra's prior research, these 2,179 motifs are categorized into 282 motif clusters, a classification determined by PWM similarity. By using this established clustering definition, nucleotide-level motif matches that are redundant are eliminated, retaining only the match with the highest score within overlapping matches belonging to the same motif cluster.

Subsequently, the scores of all non-overlapping motif matches within each motif cluster are summed, yielding one cumulative score for each of the 282 clusters. As a final step, motif binding scores for all regions within a given cell type are determined and subjected to min-max normalization across regions. This normalization facilitates model generalization and the training process, ensuring that each motif cluster's score is processed in a standardized manner.

The annotation of the 213 cell types used in pretraining followed the original cell type classification provided in the fetal and adult accessibility atlases[1–3]. This classification was achieved through clustering of ATAC-seq count profiles, with subsequent labeling based on the expression of known marker genes. The comprehensive list of tissues and cell types, along with their annotations, can be found at the Human Cell Atlas. The original atlas included 222 fetal and adult cell types, but was further filtered to remove cell types with low sequencing coverage (number of cells < 600). This approach ensured that our model was trained across diverse cellular contexts while ensuring enough coverage in the chromatin accessibility pseudobulk tracks. The data is not further balanced or curated.

###### *Input data*

GET is designed to capture the interaction between different regions and regulators. To facilitate this, we need each input sample to contain a certain number of consecutive accessible regions, mimicking the "reception field" of an RNA Polymerase II. Through previous experiments, we found that ideally the equivalent genome coverage should be around or more than 2 Mbp, a range where most of the chromatin contact happens. As a result, based on our current data prepossessing pipeline we chose to use 200 as the input region numbers for one training sample. We acquired non-overlapping samples from the genome to use as our pretraining input, and sampled from only the training chromosomes to use as our finetuning input.

For the analyses conducted in our study, we employed a fixed number of 200 peaks per sample as our standard input configuration. This number was selected based on a balance between computational efficiency and the need to encompass a representative sample of the regulatory landscape for each cell type. It's important to note that the actual genomic span covered by these 200 peaks can vary depending on several factors, including cell-type-specific variations in chromatin accessibility, the threshold applied during peak calling, and the chromosomal distribution of the peaks.

In the context of the uniformly processed datasets covering fetal and adult cell types, we have observed that 200 peaks typically correspond to a genomic range of approximately 2 to 4 million base pairs (Mbp). This estimation is derived from the understanding that the human genome, with its roughly 3 billion base pairs, yields about 150,000 accessible peaks when analyzed comprehensively. Therefore, a subset of 200 peaks would, on average, represent a genomic span of 2 to 4 Mbp, given the distribution and density of peaks across different cell types.

In terms of sampling of regions, we used non-overlapping sampling during the pretraining and a sliding window approach during finetuning. The stride of the sliding window is set to half of the number of regions in one sample (i.e. 100 peaks per step for samples with 200 peaks).

When training our main model, the borders were entirely dependent on the boundary of sampled peaks; no other priors were used. Sampling was performed independently in each chromosome starting from the beginning of the chromosomes. We have experimented with using a TAD insulation-based sampling strategy but this did not lead to significant difference in prediction performance. When doing a model interpretation for a specific gene, using a TAD-based sampling strategy for inference could incorporate useful biological priors by removing peaks unlikely to be interacting with that gene.

##### RNA-seq data processing

###### *Cell type matching*

For experiments encompassing multiomics, the correspondence between accessibility and expression is inherently determined through cell barcodes. In pseudobulk cases, where accessibility and expression are assessed independently, cell type annotations are utilized to facilitate the mapping. Specifically, the fetal expression atlas from Cao et al.[8] is employed for fetal cell types, while adult data is extracted from Tabula Sapiens[9]. When several ATAC pseudobulk share the same cell type annotation, identical expression labels are assigned. While a compromise, this approach is necessitated by the current dearth of multiome sequencing data, a situation expected to change dramatically in the near future.

###### *Expression values*

Expression values are allocated to each region within our input. Limited by poly-A scRNA-seq, only aggregated mRNA levels can be captured, resulting in values that are not reflective of the nascent transcription rate more closely tied to regulatory events. Nonetheless, these values furnish valuable cell-type-specific information. The process begins by intersecting the input region list with Gencode V40 transcript annotation to pinpoint promoters, followed by the assignment of log count per million values to regions corresponding to these promoters. All remaining regions are assigned a value of 0. Although this does not perfectly represent all transcription events happening in a cell, we believe the zero label on the non-promoter region helps in delivering informative negative labels to the model.

###### *Input target*

In alignment with the $200\times283$ input matrix, the target input is a $200\times2$ matrix, symbolizing the transcription levels of the corresponding 200 regions across both positive and negative strands.

##### Model architecture

The GET architecture consists of three parts: 1) A regulatory element (RE) embedding layer, 2) RE-wise attention layers, and 3) a linear layer as the expression prediction head, or alternatively other output head (Supplementary Figure 1).

Our GET takes 200 regulatory elements, each with 282 motif binding scores and optionally one accessibility score as a sample as the input. As a result, the input is a $200\times283$ matrix. When we choose to not use the quantitative accessibility score, we set the 283th column to 1.

We feed the sample into the RE embedding layer to generate the regulatory element embedding with a dimension of 768 for each peak. Since we do not want to lose information in the input of the original regulatory element, we apply a linear layer to capture the general information in the different classes of transcription factor binding sites. To learn the cis- and trans-interactions between regulatory elements and transcription factors, we apply 12 RE-wise Attention (REA) layers with a multi-head attention mechanism on the RE embeddings along the regulatory element.

Suppose $N_h, d_v, d_k$ denote the number of heads, the depth of values, and the depth of keys. The output from each head $h$ is computed as

$$O_h = \text{softmax}\left(\frac{X'W_q(X'W_k)^T}{\sqrt{d_k}}\right)(X'W_v), \tag{1}$$

where $W_q, W_k \in \mathbb{R}^{(n\times D)\times d_k}, W_v \in \mathbb{R}^{(n\times D)\times d_v}$ are learnable linear transformations.

Then we concatenated the output from each head $h$ for the RE-wise Attention block. The Layer Normalization (LN), Feed-forward Network (FFN), and Residual Connections are finally utilized to generate the output for each layer. Thus, the mechanism behind the RE-wise attention block is summarized as:

$$\mathbf{z}'_l = \text{MHA}(\text{LN}(\mathbf{z}_{l-1})) + \mathbf{z}_{l-1}; \mathbf{z}_l = \text{FFN}(\text{LN}(\mathbf{z}'_l)) + \mathbf{z}'_l \tag{2}$$

where $\mathbf{z}'_l, \mathbf{z}_{l-1}$ denote the intermediate representation in the block $l$ and the output from the block $l-1$. We apply two linear layers with a GELU[10] activation layer in the FFN layer.

The GET architecture is similar to the state-of-the-art model Enformer[11]. However, the following changes helped us improve and exceed its performance: GET uses the regulatory element (RE) embedding layer to capture the general information of regulatory elements in the different classes of transcription factor binding sites. Moreover, a masked regulatory element mechanism was utilized to learn the general cis- and trans-interactions between regulatory elements and transcription factors from different kinds of human cell types.

Specifically, a random set of positions was uniformly selected to mask out

$$\mathbf{M} = \{\mathbf{m}_i\}_{i=1}^{k}$$

with a mask ratio of

$$r = k/n, r = 0.5$$

. Similar to ViT-based MAE[12], We replaced the regions in the selected positions with a learnable [MASK] token, and the masked input regulatory element is denoted as $X^{\text{masked}} = (X, \mathbf{M}, [\texttt{MASK}])$, where $X = \{\mathbf{x}_i\}_{i=1}^{n}$ is the input sample with $n$ regulatory elements. The training goal is to predict the original values of the masked elements $\mathbf{M}$. Specifically, we take masked regulatory element embeddings $X^{\text{masked}}$ as input to our GET, while a simple linear layer is appended as the prediction head. Therefore, the overall objective of self-supervised training is formulated as:

$$\mathcal{L} = \mathbb{E}\left(\sum_{i\in\mathbf{M}} -\log p(\mathbf{x}_i|X^{\text{masked}})\right) \tag{3}$$

where $\mathbf{x}_i$ denotes the masked region to be predicted.

#### Training scheme

We conduct pretraining in the large-scale single-cell chromatin accessibility data. Then we finetune the pretrained model on the paired chromatin accessibility-gene expression data with the same Poisson negative log-likelihood loss function as Enformer[11]. Expression values are represented as transcript per million (TPM). We then match the cell types between RNA and ATAC datasets by annotating cell type names and ignoring those that cannot be matched. To improve training stability, we log-transform the expression values as $\log_{10}$(TPM+1). To overcome the problem that most scRNA-seq quantification is at gene-level, not transcript level, we map the gene expression to accessible regions using the following approach: if a region overlaps with a gene's transcription start site (TSS), the gene's expression value is assigned to that region as a label; if a region overlaps with multiple genes' TSS, the expression values of the corresponding genes are summed and used as the label of that region; if a region does not overlap with any TSS, the corresponding expression label is set to 0. Further, if a promoter has very low accessibility (e.g. accessibility count per million < 0.05), we also set the corresponding expression value to 0. Finally, each regulatory element is assigned to an expression target value.

The GET implementation is based on the PyTorch[13] framework. For the first training stage, we applied AdamW[14] as our optimizer with a weight decay of 0.05 and a batch size of 256. The model was trained for 800 epochs with 40 warmup epochs for linear learning rate scaling. We set the maximum learning rate to 1.5e-4. The training usually takes around a week for a cluster with 16 V100 GPUs. For the second finetuning stage, we used AdamW[14] as our optimizer with a weight decay of 0.05 and a batch size of 256. The model was trained for 100 epochs, which completes in around 8 hours using 8 A100 GPUs. Inference for all genes in a single cell type only takes several minutes, making it possible to perform large-scale screening.

#### Training details

We included a more detailed description of the optimization hyperparameters, compute infrastructure, and convergence criteria employed in the development of the model in the section below.

##### *Pretraining phase*

1. Compute Infrastructure: The pretraining of our model was conducted using 16 NVIDIA V100 GPUs or 8 A100 GPUs, reflecting the computational demands of our training dataset and the complexity of the model architecture.

2. Epochs and Duration: The model underwent 800 epochs of training, which spanned approximately one week. This extensive training period was essential for the model to learn the regulatory grammar from chromatin accessibility data across a wide array of human cell types.

3. Learning Rate: A base learning rate of 1e-3 together with cosine scheduler and linear annealing warmup in the first epoch.

4. Mask Ratio: 0.5.

5. Optimizer: AdamW with weight decay of 0.05.

##### *Finetuning phase*

1. Compute Infrastructure: Similar to the pretraining phase, finetuning was performed on 8 NVIDIA A100 GPUs, ensuring consistency in computational resources.

2. Epochs and Duration: The finetuning process was shorter, consisting of 100 epochs, and completed in around one day. This phase was crucial for adapting the pretrained model to specific gene expression prediction tasks.

3. Learning Rate: A base learning rate of 1e-3 together with cosine scheduler and linear annealing warmup in the first epoch.

4. Optimizer: AdamW with weight decay of 0.05.

5. Early Stopping: To optimize performance and prevent overfitting, we employed early stopping based on validation loss, allowing us to select the best model checkpoints for subsequent evaluation.

##### *Parameter-effecient finetuning*

GET provides the option to do parameter-efficient finetuning over any specific layer through low-rank adaptation (LoRA)[15]. A common usage of this is to adapt to a new assay or platform, in which we apply LoRA to the region embedding and encoder layers, while doing full finetuning on the prediction head. This dramatically reduces 99% parameters.

##### *Computational cost comparison*

Here we provide training batch size, RAM per batch (GPU), inference time per 1000 samples, number of model parameters, and maximum sequence length for GET, Enformer, and HyenaDNA as a measure of computational cost. We trained or finetuned GET, Enformer, and HyenaDNA for different tasks with various dataset sizes and approaches (i.e. masked pretraining on chromatin accessibility and expression head finetuning for GET; linear probing on expression for Enformer; multiclass finetuning on chromatin profile prediction for HyenaDNA). Hence we do not report a side-by-side comparison of training times for the methods.

| | GET | Enformer | HyenaDNA |
|---|---|---|---|
| Training batch size | 64 examples per batch | 12 examples per batch | 1 example per batch |
| RAM per batch (GPU) | 3 GB on 1 A6000 (200 region setting)<br>10 GB on 1 A6000 (900 region setting) | 48 GB on 1 A6000 | 35 GB on 1 A6000 |
| Inference time per 1000 samples | 2.6 seconds on 1 RTX 3090 (200 region setting)<br>13.8 seconds on 1 RTX 3090 (900 region setting) | 107 seconds on 1 RTX 3090 | 12,000 seconds on 1 A6000† |
| Number of model parameters | 86M | 251M | 6.6M |
| Maximum sequence length | Dependent on density of peak calling (typical sequence length: 2.7M)‡ | 393,216 | 1M |

**Table 1.** Comparison of GET, Enformer, and HyenaDNA models

† 1 RTX 3090 with 24 GB GPU memory was sufficient for running inference for GET and Enformer. Running inference for HyenaDNA required more RAM, in which we opted to run inference on 1 A6000 with 48 GB GPU memory.

‡ GET's effective sequence length depends on the density of peak calling for a particular region, i.e. a very sparsely sampled peak set achieves a theoretical upper bound of capturing the entire genome with one input sequence. We estimate a typical ATAC-seq peak set to contain 100,000 peaks. Assuming the peaks are evenly distributed across the genome, a 200-peak input sample into the model spans 6M base pairs, while a 900-peak input sample spans 27M base pairs. In our study, we primarily use 900-peak input samples in tandem with an atlas-level joint peak set (1M peaks), such that one input sequence encodes 2.7M base pairs on average. Capturing a receptive field at this scale aligns well with the biological prior that further genomic distances are rarely found to interact in three-dimensions, as measured by Hi-C data.

#### Model evaluation

##### *Cross-cell-type prediction*

We validated the cross-cell-type prediction performance beyond astrocytes to include a broader range of cell types. The benchmark was performed on fetal cell types with a variable length peak set defined in the original fetal accessibility atlas[4]. This comparison includes quantitative ATAC GET (n=3), binarized ATAC GET, linear probing of the Enformer CAGE output tracks trained on the Basenji[16,17] training set and inferred genes in the Basenji test set, and a training cell type mean expression baseline (**Supplementary Figure 2b**). We use Pearson correlation, Spearman correlation, and $R^2$ to evaluate the prediction performance in all settings.

To evaluation whether GET prediction preserves cell-type-specific gene expression, we compare the observed and predicted log fold change between two cell types across both seen and unseen cell types (**Supplementary Figure 2f**).

##### *Benchmarking against supervised approaches*

We have implemented comparisons with the following methods on the task of expression prediction when leaving out chromosome 11 and leaving out astrocytes, using the same input data as GET. We provide parameters used in our implementation:

1. MLP: 3 linear layers separated by ReLU (layer dimensions: 283 input, 512, 256, 2 output). SoftPlus is used for output activation.

2. CNN: 3 Conv1d layers (layer dimensions: 283 input, 128, 64, 32, 3 kernel size) followed by FC(32, 512) → ReLU → FC(512, 2). SoftPlus is used for output activation. We used the same optimizer and parameters as used in GET (base learning rate: 1e-3, cosine scheduler, linear annealing warmup, AdamW optimizer with weight decay of 0.05).

3. CatBoost: We used CatBoostRegressor with loss function "MultiRMSE" for 1000 iterations (learning rate 1e-3).

4. SVM: We used scikit-learn Support Vector Regression (SVR) with epsilon 0.2, linear kernel, and max iterations 1000. MultiOutputRegressor was used to handle 2-dimensional output.

5. Random Forest: We used scikit-learn Random Forest Regressor with 10 estimators and max depth 10. MultiOutputRegressor was used to handle 2-dimensional output.

6. Linear Regression: We opted to use linear regression instead of logistic regression because our setting aligns better with regression rather than classification. We used scikit-learn LinearRegression and MultiOutputRegressor with default parameters.

##### *Leave-out-chromosome evaluation*

We have performed the benchmark across all chromosomes and found that the performance remains consistent across chromosomes, conditioned on the same sequencing platform and data sources. We find an average Pearson correlation of 0.78 (min: 0.73, max: 0.84) on fetal astrocytes.

We also extended our evaluation of leave-out chromosomes to tumor cells from IDH1 wild-type glioblastoma (GBM) patients from the Human Tumor Atlas Network. We performed finetuning of the base GET model on tumor cells from a single patient (Case ID: C3L-03405) and evaluated performance on each leave-out chromosome. This evaluation shows an average Pearson correlation of 0.75 (min: 0.68, max: 0.81) on leave-out chromosomes.

For K562 OmniATAC prediction, we performed leave-one-chromosome-out prediction for all 22 autosomes, finding an average Pearson correlation of 0.81 (min: 0.72, max: 0.84).

For K562 CAGE prediction, we apply GET to predict K562 CAGE (FANTOM5 sample ID: CNhs12336). We first note that this comparison privileges Enformer, which was trained extensively on CAGE tracks, including K562 (track ID: 4828 and 5111), while GET must be transferred to the new assay. Here we evaluated finetuned GET against Enformer predictions summed across the two CAGE output tracks for a leave-out peak set across chromosome 14. We select chromosome 14 because it did not appear in the public Enformer checkpoint's training or validation set. Pretrained GET was finetuned in three ways:

- BATAC LoRA from BATAC pretrain: In this setting, the base model was trained on the fetal and adult atlases with binarized ATAC signal. In the finetuning, the ATAC data was binarized.
- QATAC LoRA from BATAC pretrain: In this setting, the base model was trained on the fetal and adult atlases with binarized ATAC signal. In the finetuning, we used the original accessibility CPM (aCPM) for ATAC signal.
- QATAC LoRA from QATAC finetuned: In this setting, the base model was the leave-out astrocyte RNA-seq prediction model trained on the fetal accessibility and expression atlas. In the finetuning, we used the original accessibility CPM (aCPM) for ATAC signal.

We note that these experiments leverage Low-Rank Adaptation parameter efficient finetuning to achieve significant gains in time and storage complexity. On a single RTX 3090 GPU, all finetuning converged within 30 minutes, resulting in a 3 MB K562-CAGE-specific adaptor that can be merged into the base model.

##### *Leave-out-motif evaluation*

To explore the impact of omitting motifs in the input features, we used K562 scATAC-seq data from ENCODE (accession: ENCFF998SLH) and evaluated the ATAC prediction performance when holding out randomly selected motifs. We first called peaks with MACS2 with a threshold of $q = 0.05$. We merged this peak set with the union peak set from the fetal pretraining data, keeping the peaks with at least 10 counts in K562. For finetuning computational efficiency, we used LoRA parameter-efficient finetuning of the binary ATAC checkpoint (pretrained checkpoint used for motif analysis in **Figure 4** and onward), pretrained on fetal and adult ATAC data with a 200-region receptive field.

We explored holding out randomly selected 1, 2, 3, 4, 10, and 20 motifs. For each motif, we checked whether a peak's binding score is larger than the top 20% scores in its score distribution across the genome. During the training stage, if a peak has any of the leave-out motifs passing this threshold, we set all input motif features of that peak as well as the observation accessibility count per million (aCPM) to zero. In this approach, these "knock-out" peaks do not contribute to the loss. During the evaluation stage, we calculated Pearson and Spearman correlation of aCPM only on these "knock-out" peaks with the original observed aCPM. For example, when there is only one leave-out motif CTCF, we will in effect be training with about top 20% of peaks that have low CTCF binding score on the training chromosomes, assuming evenly distributed binding sites across chromosomes. Similarly, when evaluating, we will be evaluating with 20% of peaks with higher CTCF binding in the test chromosomes. In these experiments, we evaluated on held-out chromosome 14.

In general, GET shows robust performance when leaving out 1 to 10 motifs. The performance degrades heavily when using 20 motifs with top 20% cutoff for each motif independently, due to removal of most of the training data.

##### *Platform transfer prediction*

In order to transfer to a new sequencing platform, there are a multitude of domain shifts that need to be addressed. This includes but is not limited to:

1. Sequencing depth: Lower depth will lead to fewer captured peaks. It will also affect the signal-to-noise ratio in the accessibility quantification.

2. Peak calling threshold and software.

3. Technical bias due to different library constructing and sequencing methods.

4. Biological differences.

Due to these biases, it is difficult to directly apply a model trained on one dataset to a new platform without finetuning. Thus, for a new dataset with multiple cell types available, we took a leave-out cell type approach to finetuning. For a dataset of sorted cell types where only one cell type is available, we used leave-out chromosome training.

##### *Transferring to new datasets*

The primary challenge in adapting our model to new data lies in ensuring compatibility between the input spaces of the training and new datasets. Variations in cell types, sequencing technologies, and preprocessing pipelines can result in substantially different ATAC peak sets, potentially leading to incompatible input and embedding spaces. To address this, we developed a strategy to create a compatible peak set by combining new and training peak sets. When overlaps occur between training and new peaks, we prioritize the training peak set coordinates. Unique peaks from the new data are incorporated as-is. We employed a uniform peak calling pipeline to maintain consistent peak lengths (e.g. 400 bp in the fetal-adult atlas) across training and new datasets. The comprehensive coverage of our fetal-only/fetal-adult peak set (1.3M peaks) typically results in new, unseen peaks contributing less than 10% of the total peaks. This approach has demonstrated promising transferability to various data types, including SHARE-seq data of perturbed hESC and 10x multiome GBM data.

For example, we tested a "one-shot" finetuning procedure using a single patient sample from a new dataset of glioblastoma patients. We then assessed the performance of this finetuned model against the pretrained "zero-shot" model on 16 held-out patient samples. To ensure robust evaluation, we excluded two patients from this analysis to serve as a separate test set for assessing finetuning stability. The results were promising: finetuning on a single tumor patient sample enabled GET to achieve a Pearson correlation exceeding 0.9 when predicting expression for held-out patients, while zero-shot reaches a 0.67 Pearson correlation. This demonstrates the model's strong generalization capabilities and its potential for rapid adaptation to new datasets with minimal additional training. As the availability of ATAC-seq and multiome data continues to grow, more comprehensive reference peak sets, such as the ENCODE DHS index[6] and cPeaks[18], will further facilitate the adaptation of the GET model to an even broader range of cell types and experimental conditions.

##### *Transferring to new assays*

Here we show results for transferring pre-trained GET to different functional genomics assays. For K562 bulk ATAC prediction, we collected ENCODE OmniATAC-seq data for K562 (ENCSR483RKN). After calling peaks using MACS2 with default parameters, we computed the log accessibility CPM by counting Tn5 insertions located inside the peak and filtered out the peaks with log aCPM smaller than 0.03. The remaining peaks and corresponding aCPM are used for motif scanning and prediction. We performed leave-one-chromosome-out finetuning using 200 peaks per input sample. The base checkpoint was trained on the fetal and adult atlas with the binarized ATAC setting and 200s peak per input sample. LoRA was used for all layers. Each finetuning took around 160 seconds to complete 8 epochs, when the model started to overfit. Pearson correlation was collected at 8 epochs for all finetuning. For CAGE prediction, we collected K562 CAGE (CNhs12336) BAM file from FANTOM5 and used bedtools to extract alignment counts in peaks called from ENCODE K562 scATAC-seq data (ENCFF998SLH). Finetuning was performed using 200 peaks per input sample in three settings depending on how ATAC information was used in conjunction with motif features, plus the base model used for finetuning:

1. BATAC from BATAC pretrain: In this setting, the base model was trained on the fetal/adult atlas with binarized ATAC signal, and in the finetuning we used binarized ATAC.

2. QATAC from BATAC pretrain: In this setting, the base model was trained on the fetal/adult atlas with binarized ATAC signal, and in the finetuning we used the original aCPM ATAC signal.

3. QATAC from QATAC finetuned: In this setting, the base model was the leave-out-astrocyte RNA-seq prediction model trained on the fetal accessibility and expression atlas. We further finetuned this model using quantitative ATAC signal.

##### *Dataset considerations*

Overall this suggests that certain intrinsic cellular characteristics may contribute to the observed variations in model performance. We demonstrate that GET can be applied and extended to non-physiological cell types and states and capture cell type specific transcription information. Beyond the intrinsic biological differences between cell types, we believe the following factors could also affect the performance when generalizing to new models.

1. Cell type rarity and library size (see **Figure 1E**): Rare cell types often have smaller data libraries, which can limit the model's learning potential and affect the accuracy of predictions.

2. Cell type purity and heterogeneity (comparing the perturbed hESC vs. GBM result): The dynamic and heterogeneous nature of certain cell types, such as stem cells, and the precision in identifying and classifying cell types can introduce variability in gene expression profiles, complicating the prediction task.

#### Model interpretation

In our study, we conducted thorough model interpretation analyses to ensure that GET learns useful regulatory information and offers valuable biological insights. Below, we outline the methodology employed to interpret GET.

##### *Model used for interpretation*

We trained two GET models on data with or without quantitative accessibility:

- Quantitative-ATAC Model: Accessibility scores are set to the Log10 count per million of Tn5 insertion in the given accessible region.

- Binary-ATAC Model: Accessibility scores for all regions are set to 1, focusing solely on chromatin accessibility peaks.

In our analysis and regulatory interpretation, we primarily employed the Binary-ATAC model. This approach offers improved attribution to sequence features, ensuring that the model does not overly depend on accessibility signal strength as a surrogate for sequence characteristics.

##### *Feature attribution methods*

We used multiple feature attribution methods in different analyses and provided all options to users in our packages. More specifically:

The **gradient** of the model's output with respect to the input features, represented by the vector $\nabla f(\mathbf{x})$, measures how much the model output (Expression) will change when we change a small amount of the input along a dimension (e.g. a certain motif in a cis-regulatory region). The generalization to multiple outputs in the context of neural network feature attribution extends to the Jacobian matrix $\mathbf{J}_{i,j} = \frac{\partial f_i}{\partial x_j}$,, where $f_i$ is the $i$-th output, representing the transcription level on either the positive or negative strand, and $x_j$ is the $j$-th input feature, comprising scanned and summarized binding scores for 282 TF motif clusters, and an additional dimension for accessibility scores. This formulation enables the computation of the Jacobian matrix, vital for understanding the influence of individual features on the transcription levels.

##### *Enhancer-gene pair prediction*

We restrict the benchmark dataset to either the fetal erythroblast peak set or K562 DNase peak set for a fair comparison. To get the enhancer importance score for each gene from GET, we used the $\ell^2$-norm of the region embedding layer Jacobian and weighted it with aCPM of each region as the GET Jacobian score. We note that this procedure could potentially be improved in the future: for example, to use random genomic background as the baseline for the Jacobian calculation, and other interpretation methods like Integrated-gradients[19] or DeepLift [20] could potentially be used. However, we believe current benchmark dataset size for this task is still limiting comparing to the genome scale ($10^4$ measured pairs v.s. $10^6$ to $10^7$ required measurements of genome wide E-P interaction). Thus we leave the systematic optimization for this task to future study. For other scores used in this study:

1. ABC: We computed ABC Powerlaw by multiplying the powerlaw function in the official ABC repo with $\gamma = 1.024238616787792$ and scale $= 5.9594510043736655$, values which were trained on K562 Hi-C data and provided in the same repo.

2. Enformer: For Enformer, we used Enformer's contribution score (gradient × input) with background normalization, following the normalization procedure described in Gschwind et al[21].

3. HyenaDNA: For HyenaDNA, we used the largest pretrained model available through Hugging Face (context length 1 million base pairs). To score enhancer-gene pairs, we performed *in silico* mutagenesis (ISM) by knocking down the enhancer element (i.e. setting each base pair in the enhancer region to the unknown nucleotide "N" in the vocabulary set) and comparing against wild-type likelihood of observing the promoter sequence.

4. DeepSEA: Nucleotide-level DeepSEA results were retrieved directly from the original publication and averaged over each peak.

All score in this benchmark (ABC, Enformer, GET, HyenaDNA, DeepSEA and DNase/ATAC) are further normalized across each gene's $\pm 100$ peaks to make them comparable across genes.

Recent studies have highlighted the dominant importance of 1D genomic distance in governing CRISPRi enhancer knockout effects (e.g. Gschwind et al.[21]). In this benchmark, most methods include a component of genomic distance. For example, Enformer incorporates exponential decay in its positional encodings. HyenaDNA incorporates a sinusoidal positional encoding over the DNA sequence, and our benchmarking results follow an exponential decay from the TSS (see **Figure 3c** NFIX). We have also extended GET to incorporate distance information. In particular, we designed a simple DistanceContactMap module for GET to convert the pairwise 1D distance map between peaks to a pseudo-Hi-C contact map. DistanceContactMap is a simple 3-layer 2D convolutional neural network (kernel size 3) with $\log_{10}(\text{Pairwise Distance} + 1)$ as input and SCALE-normalized observed contact frequency as output. A Poisson negative log-likelihood loss was used to train the model. We trained DistanceContactMap with the same K562 Hi-C data (ENCFF621AIY) used for training ABC Powerlaw, resulting in a 0.855 Pearson correlation, which mostly captured the exponential decay in contact frequency. We termed the prediction of this model "GET Powerlaw." The other two scores shown in Figure 3d are defined as:

1. GET (Jacobian, DNAse/ATAC, Powerlaw) = GET Jacobian + aCPM × GET Powerlaw
2. GET (Jacobian, Powerlaw) = GET Jacobian × GET Powerlaw

This model can be further improved as future work by taking GET region embeddings as additional input and learning to predict cell type specific 3D contacts.

###### ***LentiMPRA zeroshot prediction***

The experimental procedure involves designing a library of lentivirus vectors that contain both desired sequence elements and a minipromoter. The vector will be randomly inserted into the genome through viral infection; the regulatory activity is then measured through sequencing and counting the log copy number of transcribed RNAs and integrated DNA copies.

To simulate this approach using GET, we first collected the sequence element library and constructed the vector sequence for each mini promoter. We then follow the same data preprocessing procedure to get the motif score of the inserted elements. For each element, we perform *in silico* insertion by summing up its motif score with an existing region on the genome. The $\pm 100$ regions centered around the insertion region were then used as an input sample for GET to make expression prediction. The mean predicted expression $\log_{10}$ TPM was multiplied with the mean predicted accessibility as the predicted regulatory activity. For each region, we perform 600 insertions across the genome to match the experimental insertion count. We used the GET model finetuned on K562 NEAT-seq data to perform the inference. In total, *in silico* lentiMPRA of all 200,000 elements in K562 took around 5 days to finish.

For Enformer, we performed the same analysis, with the only difference being that we integrated the vector sequence to a random position on the genome and collected a 196,608 bp sequence centered around the insertion site. Enformer is trained on 5,313 human epigenome tracks, with 486 experiments specifically for K562. To compute the regulatory activity, we selected the output from the K562 CAGE track, which is a quantitative and nucleotide-level map of 5' of transcripts. Following the practice of the original study, we used the average output of the 3 bins in the center of the sequence as the predicted expression for a sample. Each element was also inserted into 600 random genome locations to compute the final averaged regulatory activity. We were only able to perform these experiments for 1,000 enhancers and 1,000 non-enhancer elements due to the time complexity of Enformer inference. The comparison with GET is performed on the same set of elements.

We stratified the K562 lentiMPRA elements (approximately 200,000) by overlapping the annotated 15 ENCODE ChromHMM states computed from histone mark and other ChIP-seq data for K562. We selected the elements overlapping with states "12 EnhBiv," "6 EnhG", and "7 Enh" as enhancers, and "13 ReprPC," "14 ReprPCWk," and "15 Quies" as repressive and quiescence regions.

In our manuscript, we followed the categorization of the different types of elements originally provided by Agarwal et al.[22]. "Promoters" are defined as protein-coding gene promoters, "Peaks" are defined as accessible peaks called from ENCODE K562 ATAC-seq data (which contain most of the enhancers), "Heterochromatin" are defined as genomic tiles of heterochromatin regions around selected genes (e.g. GATA1), and "Control Group" are defined as synthetic elements.

Using all K562 (untreated) ChIP-seq peaks from the ChIP-Atlas, including 32 targeting histone marks and 562 targeting TFs (more than 500) and other targets (e.g. G-quadruplex, AGO1/2, Cas9), we performed enrichment analysis against the lentiMPRA promoter (approximately 57,000) and ATAC peak (approximately 169,000) library, which was further subdivided into 4 groups respectively based on GET prediction (P) (P+: >1; P-: <1) and observed readout (O) (O+: > 0.5; O-: <0.5, visualized in the Supplementary Figure 3a).

The enrichment Fisher's exact test was performed using the bedtools fisher command. The overlapping ChIP-seq peaks for the same target were merged before testing. Log2 fold enrichment for significant results ($p < 0.05$ in any group) were visualized and compared using a heatmap for all histone marks (Supplementary Figure d) and the 100 most variable TFs in terms of enrichment across the four expression-based groups (Supplementary Figure e). We noticed that O+ promoters showed larger H3K4me3 enrichment, and P+ promoters showed larger H3K4me3 and H3K27ac enrichment. Interestingly, P+O- promoters have a distinct signature of H3K9me3, H2AK119Ub, and H3K79me2 compared to the P+O+ group, potentially relevant to promoters that were originally regulated by the Polycomb complex or in heterochromatin. Since lentiMPRA elements are relatively newly inserted to the genome, these repressive mechanisms may have not yet been set up, leading to a discrepancy between RNA-seq based prediction and lentiMPRA readout. Similarly, accessible peaks are enriched in H3K27ac and¬ H3K4me1/2/3 in general, while P+O+ peaks are particularly enriched in H3K122ac, a signature of active enhancers and transcription36. Comparing P-O+ peaks to P+O+ peaks, H3K27me2, H3K14cr, and H4K20me1 are more enriched. Pioneer factors like GABPA, NFYA, and NEUROD1 are enriched in P+ and particularly P+O+ promoter and enhancers. Other signature transcription factors are CEBPZ (P+ vs. P- promoters and P-O+ vs. P-O- peaks) and WDR5 (P+O+ vs. P-O+ peaks).

##### Identifying important regions and regulators

We first gather inference samples across the genome by producing 200-region windows that centered around each gene's promoter. Given a specific gene $g$ on strand $s \in \{0,1\}$, the expression value can be inferred using the GET model $f$ applied to an input matrix $\mathbf{X} \in \mathbb{R}^{r\times m}$, where $r$ denotes the number of regions, and $m$ includes motifs and optionally accessibility features:

$$\mathbf{E} = f(\mathbf{X}) \tag{4}$$
$$E_g = \mathbf{E}[r//2, s] \tag{5}$$

where $[\cdot,\cdot]$ is the indexing operator and s is the strand of the gene.

The Jacobian matrix (tensor) $\mathbf{J}_X \in R^{r\times 2\times r\times m}$ of $f$ at the point $(\mathbf{E},\mathbf{X})$ evaluates how each output dimension will change when each input dimension changes a small quantity. We specifically pick the output dimension and strand that correspond to the given gene, represented as $\nabla g \in \mathbb{R}^{r\times m}$:

$$\nabla\mathbf{g} = \mathbf{J}_X[r//2, s] \tag{6}$$
$$\mathbf{J}_\mathbf{X} = \frac{\partial \mathbf{E}}{\partial \mathbf{X}} \tag{7}$$

The feature (motif) importance vector $v_g \in \mathbb{R}^m$ is obtained by multiplying the gradient element-wise with the original input and summarizing across regions:

$$v_g = \sum_{i=1}^{r} (\nabla\mathbf{g} \odot \mathbf{X})[i,:] \tag{8}$$

where $\odot$ signifies the element-wise or Hadamard product. Since the gene-by-motif matrix is mostly used for feature-feature interaction analysis, we use the $\mathbf{X}$ with quantitative ATAC signal even when we infer $\mathbf{J}_\mathbf{X}$ using a "Binary-ATAC" model. This helps us to study the relationship between regulators and observed chromatin accessibility.

The cell type $c$ specific genome-wide gene-by-motif matrix $\mathbf{V_c}$ is acquired by concatenating the $v_g$ across the genome. The same process can be applied to different cell types.

Similarly, the region importance vector $l_g \in \mathbb{R}^r$ is given by:

$$l_g = \sum_{j=1}^{m} (\nabla\mathbf{g} \odot \mathbf{X})[:,j] \tag{9}$$

In practice, we use the jacobian of the region embedding with regard to output for calculating the region importance score as the embedding score distribution is less skewed than the input motif binding score, potentially make the jacobian more comparable across regions.

##### Gene ontology enrichment of top target genes of a regulator

Based on the gene-by-motif matrix $V_c$, we can choose a TF/motif (in our case, GATA) and ask which genes will be mostly affected by this TF by identifying the largest entries in the motif column. We chose the top 1,000 genes and performed gene ontology enrichment analysis using g:Profiler with the default "g_SCS" multiple hypothesis testing correction. To avoid general terms we filtered the result with term size (gene number in a term definition) larger than 500 and smaller than 1000. Terms with adjusted P-value smaller than 0.05 are retained as significant terms. We further selected TFs in the "Hemopoiesis" term with expression $\log_{10}$TPM > 1 for visualization against the GATA motif score.

##### *Transcription factor and target gene correlation*

In this analysis, we sought to elucidate the relationship between transcription factors (TFs) and their target gene expression across different cell types. Gene-by-motif files were aggregated and organized into a unified structure comprising genes, motifs, and corresponding cell features. We identified the target genes for each TF within predefined motif clusters and computed the mean expressions of both the target genes and the corresponding TFs. To avoid potential artifacts caused by the experimental batch effect in expression measurement, we performed the analysis both in adult and fetal cell types and also in only fetal cell types and found similar results. The analysis was performed iteratively for all TFs within the motif clusters specific to fetal cell types.

##### *Regulatory embedding*

We collected the embedding of each gene after each transformer block of GET. For a gene $g$, its embedding is defined as the embedding vector of the promoter in the output of the $i$-th block. The embedding contains not only promoter information but also information from surrounding regions owing to the attention mechanism. In general, the deeper the layer, the more its space is dominated by the expression output (**Supplementary Figure 5b**). tsne-cuda was used to visualize the embedding due to data size. Louvain clustering was performed on the embedding space to colorize the visualization. Resolution is arbitrarily chosen to keep the cluster number around 10 and close to the UMAP density. For cell-type based subsampling, UMAP[23] was used instead for visualization for better visual separation between clusters.

We computed the embedding in two different settings: the cell type specific setting, in which each dot is a gene embedding from a specific cell, and the cell type agnostic setting, in which each dot is a gene embedding randomly sampled from all cell types. 50,000 embeddings are sampled in the second case to make the UMAP computation feasible.

##### *Causal discovery of regulator interaction*

We performed pairwise Spearman correlation using the gene-by-motif matrix in both cell-type-specific and agnostic settings. Input × gradient scores were used to construct the matrix for computational efficiency. For the cell type agnostic settings, all genes with their promoter overlap with open chromatin peaks from all cell types were used in the correlation calculation. Causal discovery was performed on the gene-by-motif matrix using LiNGAM[24]. For the cell type agnostic settings, 50,000 genes were randomly sampled from all cell types, and the resulting matrix was subjected to the LiNGAM algorithm implemented in the Causal Discovery Toolbox Python package with default parameters.

To benchmark the predicted causal edges in the cell type agnostic setting, we downloaded the known physical interaction subnetwork from the STRING V11 database[25] and kept interactions with a combined score larger than 400 as the ground truth label. Since the pairs predicted by GET are on the motif cluster level, we mapped the physical interactions between TFs onto the motif clusters based on the motif cluster annotation. The resulting motif-motif physical interaction network was then compared with our prediction to calculate the precision. We also downloaded and compiled all significant interactions determined by mass spectroscopy[26] and mapped them also to motif-motif interactions for comparison. For comparison with ChIP-seq colocalization, we acquired colocalization result between ChIP-seq track for 677 transcription factors in HepG2, acquired from TF Atlas. The method for calculating colocalization is documented in the ChIP-Atlas repo. Each ChIP-seq peak set was stratified into 3 tiers (high, mid, low). Then for a pair of transcription factors P1 and P2, the authors check colocalization between every tier pair and assign scores with a preference for high-high colocalization (score=9). If the strong binding peaks of P1 overlap with strong binding peaks of P2, the P1-P2 interaction is more robust than the case in which the P1 strong binding sites only overlap with the P2 weak binding sites. We presented the colocalization stronger than mid-mid (score=3) interactions (score>=4) in **Figure 4h** as those present the more reliable interactions. A stronger cutoff (score>=9, keeping only high-high interactions) reduced its performance to 0.097 macro F1 score at 2% recall.

For comparison with motif colocalization, we collected the GET input matrix (accessible-region-by-motif) for hepatocyte or concatenated the input matrix across all fetal and adult cell types. Pairwise pearson correlation was computed across all collected regions, resulting in a score for every pair of motifs.

For the cell-type-specific motif-motif interaction in the GET catalog, we performed causal discovery using the gene-by-motif matrix for all cell types. Interactions with the top 5% absolute effect size were retained in the final database. For each interaction, we performed structural analysis between the two TFs with the highest expression in the corresponding cell types.

#### Structural analysis

##### *Alphafold benchmark on intra-family binder prediction*

We classify a TF as an intra-family binder if any of its member TFs have a known physical interaction annotated in the STRING V11 database. Based on the hypothesis that if a TF can bind as a heterodimer, it should also have the potential of binding as a homodimer due to sequence and structure similarity (although the dimerization affinity might be different). We thus used AlphaFold to predict the hypothetical homodimer structure of all known TFs and tried to predict whether a TF could be an intra-family binder based on various AlphaFold-based metrics. We used several different AlphaFold-based metrics, including mean_plddt (average predicted Local Distance Difference Test score across all residues), pAE (predicted Aligned Error across all inter-chain interactions), pDockQ (predicted DockQ metric using interface pLDDT), and pDockQ × pAE. We found that pDockQ ×pAE led to the best AUROC (0.69) and AUPR (0.41) when classifying intra-family binder TFs.

###### *Protein sequence segmentation*

pLDDT from AlphaFold is a reliable protein domain caller due to its accurate structure prediction performance. We segment each TF protein sequence into low and high pLDDT regions. Empirically, we found that 80% (recall) of known DNA-binding domains can be easily identified using high pLDDT regions plus a high ratio of positively charged residues. More specifically, we first computed smoothed pLDDT using a 10 amino acid moving-average kernel and then normalized the score by dividing the max. After that, any region that has a smoothed pLDDT score less than 0.6 is defined as a low pLDDT region. If two low pLDDT regions are close (<30 aa) they will be merged as one. Any region that is not a low pLDDT region will be labeled as a high-pLDDT region.

###### *Multimer structure prediction*

LocalColabFold and ColabFold are used to predict multimer structures with the AlphaFold Multimer v2.3 model. For homodimer prediction, we used all 5 models with 3 recycles. For our large-scale interaction screening, we used model 3 and 3 recycles for each prediction. Predicted aligned error (PAE) and predicted LDDT were stored for downstream analysis. pDockQ was calculated following code from FoldDock[27].

If the multimer structure has a newly appeared peak, we treat it as potential evidence of potential interaction. PAE, pDockQ, and ipTM were further checked to assess the confidence of the interaction. After AlphaFold 3 was released, we re-performed structure prediction for full length PAX5-NR2C2 sequences and identified the same PAX5 G183-NR domain interaction.

###### *Molecular dynamics simulation*

The initial configuration was prepared from the AlphaFold predicted PDB file. The Amber99SB-dispersion (a99SBdisp) force field was employed for system parameterization. A cubic simulation box was defined with a box size of 1 nm. Subsequently, the system was solvated using the TIP4P water model through the solvate module. To neutralize the system and generate physiological ion concentrations, sodium (Na+) and chloride (Cl-) ions were added using the genion module. The energy minimization terminates upon reaching a maximum force below 1000.0 kJ/mol/nm. Each minimization iteration utilizes a step size of 0.01 and is configured to run for a maximum of 50,000 steps. The system was then equilibrated in two steps: first in the NVT (Constant Number, Volume, Temperature) ensemble and then in the NPT (Constant Number, Pressure, Temperature) ensemble for 100 ps of simulation time. A 100-ns production run was then performed and trajectories and energy profiles were stored for subsequent analysis. All configs of these are available at the Proscope repo.

###### *Structure visualization*

ChimeraX was used to visualize the predicted structures. VMD was used to generate the movie of molecular dynamics simulation trajectory.

##### Biological experiments

###### *TFAP2A co-immunoprecipitation*

HeLa cells were purchased from ATCC (CCL-2). HeLa cells were cultured in DMEM (Gibco, 11965) supplemented with 10% defined FBS (HyClone, SH30070), at 37°C/ 5% CO2. HeLa cell protein lysates were generated with 0.5% NP-40 lysis buffer (50mM Tris-HCl, 150mM NaCl, 0.5% NP-40) with phosphatase and protease inhibitor cocktail (Sigma-Aldrich, PPC1010). Samples were incubated with 5 µg agarose-conjugated TFAP2A primary antibody (Santa Cruz Biotechnology, sc-12726 AC) overnight at 4°C. Beads were washed, then boiled in Laemmli loading buffer (BioRad, 1610737). Proteins were separated on 10% Tris-Glycine gels (ThermoFisher, XP00100), transferred to PVDF (Immobilon-P, IPVH00010) and probed with primary antibodies against TFAP2A (ABclonal, A2294), ZFX (ThermoFisher, PA5-34376) and $\beta$-ACTIN (Santa Cruz Biotechnology, sc-47778) followed by chemiluminescence detection.

###### *Proximity ligation assay to detect PAX5-NR2C2 interactions*

We initially cloned PAX5 wild type (WT) and PAX5 G183S mutant into the pCDNA3.1-MCS-13Xlinker-BioID2-HA (Addgene 80899)[28]. After verification, we subcloned PAX5-WT-13Xlinker-BioID2-HA and PAX5-G183S-13Xlinker-BioID2-HA into the pCDH-GFP-puro vector (System bioscience CD513B-1). We transduced the REH B-ALL cell line with pCDH- PAX5-WT-13Xlinker-BioID2-HA-GFP and with pCDH- PAX5-G183S-13Xlinker-BioID2-HA-GFP and selected transduced cells with puromycin (1ug mL-1) to generate stable cell lines. Proximity ligation assay (PLS) was performed following previously published methods[28–30]. Briefly, REH stable cell lines with control vector pCDH_13Xlinker-BioID2-HA-GFP, pCDH_PAX5-WT-13Xlinker-BioID2-HA-GFP and pCDH_PAX5-G183S-13Xlinker-BioID2-HA-GFP were incubated with 100uM biotin (Sigma Aldrich B4501) for 24h. We harvested the cells, washed them twice in cold PBS and incubated them for 50min on ice with occasional vortexing in lysis buffer (150 mM NaCl, 10 mM KCl, 10 mM Tris-HCl pH 8.0, 1.5 mM MgCL, 0.5% IGEPAL) supplemented with protease and phosphatase Inhibitor (Life Technologies 78443) and 63U of benzonase (Sigma Aldrich 70746-3). Proteins were clarified by centrifugation at 21,000g for 15min at 4C. We performed total protein quantification using the Pierce BCA Protein Assay kit (ThermoFisher Scientific 23225) and incubated 1mg of total protein extract with 100ul of magnetic Streptavidin beads (Dynabeads MyOne Streptavidin C1, Life Technologies 65002) on a rotator at 4C overnight to isolate biotinylated proteins. We washed the beads twice with lysis buffer, once with 1M KCL, once with 0.1M Na2C03, once with 2M Urea 10 mM Tris-HCL pH 8.0 and twice again with lysis buffer. Biotinylated proteins were eluted by boiling them in 4X protein loading buffer supplemented with 2mM biotin and 50mM DTT at 95 °C for 10 min. Biotinylated proteins in total protein extracts or immunoprecipitated were detected by western blot using standard protocols and the following antibodies: streptavidin-HRP antibody (Life Technologies S911), anti-PAX5 (Cell Signaling 8970), anti-HA (Cell Signaling 3724), anti-NR2C2 (Cell Signaling 31646), anti-NCOR1 (Cell Signaling 5948) and NRIP1-HRP (Santa Cruz Biotechnology sc-518071). Proteins were detected using a Li-Cor Odyssey OFC instrument and quantified using the GelAnalyzer 23.1 software.
